# Hebbian learning generates individuality in spatial decision-making

**DOI:** 10.64898/2026.10.12.758645

**Authors:** Mohammad Salahshour, Iain D. Couzin

## Abstract

Individuals exposed to similar environments often develop consistent behavioral differences, yet how learning generates individuality remains unclear. We study agents whose neural connections develop through unsupervised Hebbian learning during pursuit of a moving target. All agents begin with identical networks without recurrent connections and neurons tuned to world-fixed directions. The target coactivates neurons with overlapping directional receptive fields while leaving more distant neurons relatively inactive. Hebbian learning converts this activity pattern into local excitation and longer-range inhibition. Stochastic differences in target motion and neural activity during learning produce individual-specific departures from each network’s average center–surround profile. We then freeze the connections and repeatedly test agents with two equally attractive targets, a task absent from training. Individual agents tend to spend more time near one target than the other, with preferences differing among agents but remaining repeatable across trials. Replacing each network with its average connectivity profile strongly reduces preference strength and leaves little repeatability. Rearranging which connections are stronger or weaker than average, while preserving the amount of variation and average profile, retains repeatable preferences and improves tracking in a separate moving-target test. However, preferences confer no general advantage when targets differ in value. Under standard sensory and inhibitory settings, repeatable preferences and accurate choices coexist only at intermediate neural noise levels tested. A matched egocentric model, with tuning relative to the agent’s heading, shows no detectable target-side repeatability. Hebbian learning can therefore generate individual spatial preferences, whose expression depends on how the network represents space.

## I. INTRODUCTION

Animals navigate by combining sensory cues, memory, and internal state, yet individuals in the same environment often move and decide in persistently different ways. Such consistent between-individual differences are termed animal personalities, whereas correlations among behavioral traits are known as behavioral syndromes [1– 6]. Genetic and environmental variation contribute, but stochastic processes shape individuality even in their absence. Noise during neurodevelopment produces distinct neural circuitry across individuals, and with it differences in cognition and behavior [6–10]. Such variation shapes how populations respond to change and how their social structure forms [3, 11], yet how developmental noise becomes individual differences in decision-making is unresolved at a mechanistic level [7, 12, 13].

Idiosyncratic behavioral biases, such as those arising from lateralization, covary with problem-solving and cognitive performance across taxa, from birds [14–16] to primates [17] and humans [18]. What ties an individual’s biases to its cognitive performance is not known, and we do not attempt a general account here. The observation motivates a narrower question, whether one developmental process can both give a neural network an individual-specific bias and shape how it makes spatial decisions. Hebbian learning, in which the connection between two neurons that are active at the same time grows stronger, is one process that could do both, and we study it in agents built on a ring attractor network.

Continuous attractor networks are a natural place to ask this. Neurons on a ring with local excitation and longer-range inhibition sustain a bump of activity that encodes a continuous variable [19, 20], a scheme applied to head-direction, place, and grid cells [21–24]. How such connectivity arises is a separate question from how it functions. It has been prescribed by hand [22, 23, 25, 26], trained toward explicit spatial objectives [27–29], and related to the efficient representation of space [30–33]. Unsupervised plasticity builds such structure as well, with associative learning self-organizing head-direction attractor connectivity [34], spike-timing-dependent plasticity developing grid connectivity in an unstructured spiking network [35], and Hebbian learning with adaptation yielding grid fields from spatial input [36]. These studies established mechanisms for self-organizing spatial circuitry [34–36]; here we ask how such learning also generates repeatable differences among individuals.

Real attractor circuits are heterogeneous [37, 38], and because they form under the same stochastic developmental conditions that generate between-individual differences, heterogeneous connectivity and behavioral individuality plausibly share an origin [12, 39–42]. How neural heterogeneity shapes computation remains incompletely understood. In task-trained entorhinal models, virtual lesions show that heterogeneous cells contribute to function rather than degrade the code [37], and their contribution depends on whether the task is path integration or the construction of spatial functions [28]. More generally, in complex systems heterogeneity can degrade or enhance function, sometimes by extending the responsive regime near a critical point [43–46]. Hebbian learning [47, 48] is a natural candidate for a rule that builds such circuits, including their heterogeneity. It underlies self-organizing feature maps [49] and structural properties of networks from orientation selectivity [50] to continuous attractor connectivity [34, 35, 51, 52], and under stochastic conditions it can produce individual-specific departures from a shared connectivity profile.

Ring attractor networks are a fundamental component of spatial navigation in invertebrates [53, 54] and vertebrates [20, 22], and their dynamics bear on spatial choice as well. Sridhar et al. introduced a model of decisionmaking among spatial targets in which neural groups each code one target’s direction and the choice is set by the equilibrium the network settles into [55]. Coupling between groups is set by the egocentric geometry of the targets, the angle the pair subtends from the agent’s position, so the effective connectivity changes as the agent moves. Neural groups representing similar directions reinforce one another, whereas widely separated ones compete, so an animal can first move in a compromise direction between the targets; as it approaches, their angular separation grows, and at a critical separation the network shifts abruptly from compromise to one target, a transition known as a bifurcation. With more than two targets, successive transitions narrow the options until one is selected [55, 56]. Choice emerges through continuous feedback between neural activity, movement, and target geometry, and this behavior has been demonstrated in fruit flies, desert locusts, and larval zebrafish [55, 57, 58].

The model that we build on here, which we introduced in depth in our work on allocentric flocking [59] and which also appears in our study of collective motion in locusts [58], is constructed differently. Neurons are not linked to targets. Each has a preferred direction fixed either in world coordinates (allocentric) or relative to the agent’s heading (egocentric), and receives external input through a receptive field tuned to that direction, representing, for instance, an animal’s visual field (Fig. 1A). The connectivity is likewise fixed, a function of the angular distance between neurons on the ring rather than of the agent’s position in space. Every neuron combines the sensory input it receives with the recurrent input arriving from the rest of the network, analogous to models of collective information acquisition in which individuals integrate personal and social information [60]. Here the neuron corresponds to the individual, its receptive field to personal information, and the recurrent input to the social cue. This framework allowed us to compare behavior under egocentric and allocentric representations of space, including how the reference frame shapes collective movement [59]. Both forms of spatial representation occur in biological navigation [54, 61–64].

**FIG. 1.**
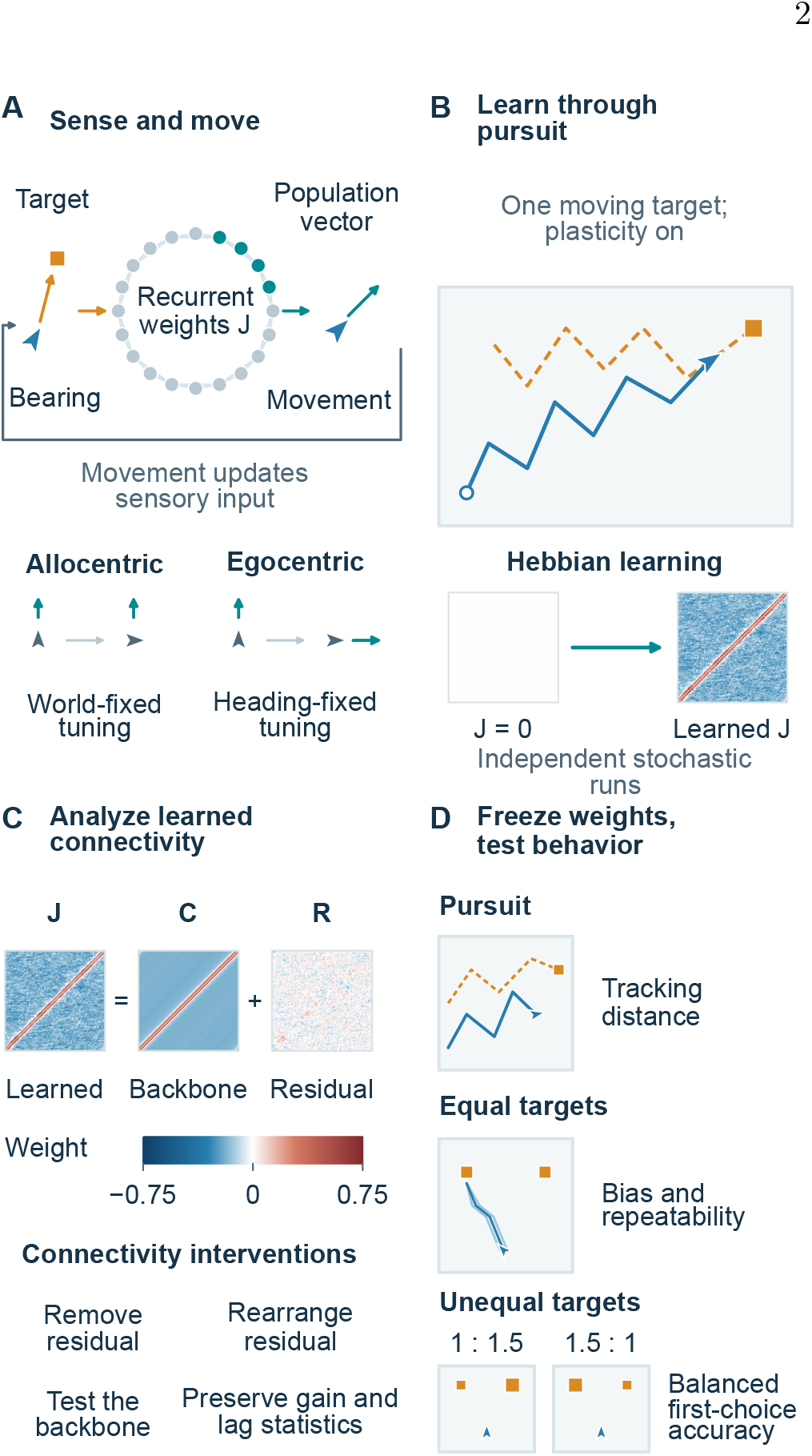
A directional neural circuit links learning to spatial behavior. (A) Target bearings drive a ring of binary neurons; teal nodes denote active neurons. Recurrent input and stochastic neural updates shape the activity, whose population vector determines movement. Preferred directions are fixed in the world in the allocentric model and relative to the agent’s heading in the egocentric variant. Both sensory input and movement use the chosen frame. (B) During initial training, networks learn while pursuing one moving target. Starting from no recurrent connections, independent stochastic runs produce distinct learned connectivities. (C) A learned connectivity *J* is decomposed into a network-specific circulant backbone *C*, shared across neurons within that network, and a residual *R*. Matched controls test the amount and arrangement of residual heterogeneity while controlling coupling gain. Matrix rows index postsynaptic neurons and columns presynaptic neurons. (D) With connectivity frozen, repeated trials measure pursuit, preference and repeatability between equal targets, and first-choice accuracy between unequal targets presented in both orientations. Continued plasticity and preformed-ring starts provide additional tests. Paths are schematic illustrations of the tasks, not simulated observations; the connectivity matrices illustrate one learned network. Numerical protocols are given in the Methods.

The model also generates spontaneous movement in the absence of targets. At high neural noise, activity fluctuates without sustaining a preferred direction, and in this disordered phase the agent follows a random walk. At lower noise it organizes into a persistent bump whose direction is anchored to the environment by the allocentric coding, giving an ordered phase in which the agent moves consistently at a spontaneously chosen bearing, along a relatively directed trajectory. We also demonstrated that when agents provide input to one another, self-organized collective motion arises [59]. A single agent tracks a moving target most closely near the order–disorder transition between these two phases, where a weakly ordered network re-orients readily but travels slowly, so that movement trades speed against accuracy [44–46, 59].

Building on this work, we explore the role of heterogeneous connectivity in ring attractor networks. In our previous studies the connections followed the same modified cosine function in every agent. Neurons representing similar directions excited one another, and as angular separation grew excitation weakened and gave way to inhibition, strongest between opposite directions. The connections were also symmetric, so each pair of neurons influenced one another equally. That work generated diverse movement patterns but did not consider how learning from experience could shape a network, differentiate individuals, or bias their choices.

Here, we allow Hebbian learning to shape the connectivity and then either freeze the learned weights, or let the same learning rule continue during behavioral testing (which we term continued plasticity). Initial training involves only single-target pursuit (Fig. 1B), so neither choice task is encountered before testing. We ask whether this experience generates a shared center–surround connectivity profile together with individual-specific deviations, and whether those deviations produce repeatable preferences between equally attractive targets. We also introduce heterogeneity directly into otherwise homogeneous networks to distinguish the effects of variation itself from those of the particular structure produced by learning. By manipulating the learned connectivity, we examine how the amount and arrangement of heterogeneity affect both pursuit and choice, including accuracy when targets differ in value (Fig. 1C). The agents perform three spatial tasks involving either one moving target or two stationary targets (Fig. 1D). Each target provides a directional cue by exciting neurons tuned to its bearing relative to the agent, without conveying its distance. The first task measures pursuit of a single target performing a random walk. The second tests whether an agent maintains commitment to one of two equally attractive stationary targets. Commitment here means sustained or repeated use of a target zone, without a stopping or reward rule at arrival. The third tests whether it selects the more attractive of two targets, represented by a stronger sensory input. Finally, we investigate how these outcomes depend on neural noise, the rate and duration of continued Hebbian learning, and the network’s initial connectivity. The main text focuses on the allocentric model. A matched egocentric comparison is reported in Supplementary Note S15 and Supplementary Fig. S1, with its principal findings summarized in Section II H 3. The model and study design are summarized in Fig. 1.

## II. RESULTS

### A. The model

We begin with the ring attractor model introduced in our work on allocentric flocking [59]. It provides a homogeneous baseline in which every agent has the same fixed pattern of connections. The model is deliberately simplified. It does not reconstruct a particular circuit, and we use it to ask which features of heterogeneity are generic. Each agent has a network of *N* neurons whose collective activity determines its movement, with *N* = 100 except where noted. Neurons are modeled as binary spins, *s*_*i*_∈ {− 1, +1}, with *s*_*i*_ = +1 denoting an active neuron and *s*_*i*_ = − 1 an inactive one. Such spin representations are standard in models of neural computation and associative memory [48, 65]. In the allocentric model, neuron *i* has a preferred direction *α*_*i*_ = 2*π*(*i* − 1)*/N* fixed in world coordinates. In the egocentric variant, these directions rotate together with the agent’s heading, so each neuron’s tuning remains fixed relative to that heading. In both cases, the same preferred directions determine sensory input and movement. A receptive field is centered on each neuron’s current preferred direction, so that a target at bearing *θ*_*t*_ excites the neuron through a circular Gaussian of standard deviation *σ* = 2*π/N*. Sensory input is thus mapped topologically onto the ring. A target drives most strongly the neuron whose preferred direction matches its bearing, and progressively less strongly the neurons to either side.

The recurrent input to neuron *i* is set by the connection weights *J*_*ij*_. In the homogeneous baseline those weights depend only on the angular separation of the two neurons, so every neuron sees the same profile of couplings, exciting neurons tuned to nearby directions and, as preferred directions grow apart, weakening and then reversing to inhibition. This is the standard center–surround organization of ring-attractor models, which we refer to as the *cosine ring*, defined by

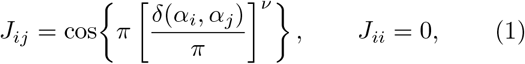

where *δ*(*α*_*i*_, *α*_*j*_) is the circular angular distance between the two preferred directions, the length of the shorter arc that separates them, so that it lies between 0 and *π*, and no neuron is coupled to itself. The tuning exponent *ν* fixes how far the excitation reaches. The coupling changes sign at an angular separation of *π*2^*−*1*/ν*^, which is 90^*°*^ for a plain cosine and 45^*°*^ for the *ν* = 0.5 used here, so the profile we use has a narrower excitatory core and a broader inhibitory surround than a plain cosine.

Recurrent and external inputs combine into an incoming local field on neuron *i*,

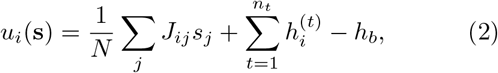

the second sum running over the *n*_*t*_ targets in view. At each neural update one neuron is selected at random and its flip is accepted with probability min {1, exp[− 2*βs*_*i*_*u*_*i*_], where *β* sets the inverse noise of the dynamics. For a symmetric connectivity this local-field rule is the single-spin Metropolis dynamics of an equilibrium Hamiltonian. The learned networks are generally asymmetric, and we treat them as nonequilibrium stochastic networks defined directly by their local fields. After *T*_0_ neural updates the agent moves according to the vector sum of its active neurons’ preferred directions, scaled by *v*_0_*/N*,

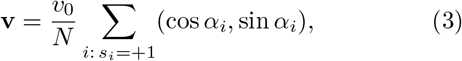

with *v*_0_ = 10, so the state of the network is continuously converted into motion. The couplings are scaled by the conventional 1*/N* factor, which keeps the energy extensive and makes *β, h*_0_, and *h*_*b*_ the physical inverse noise, stimulus amplitude and input offset, respectively. The full model, geometry, and numerical settings are given in the Methods and Supplementary Note S1.

The term *h*_*b*_ in Eq. 2 is a constant global inhibition subtracted once from each neuron’s field, independent of the number of targets. We vary it to raise or lower the net input reaching the network. Unless an offset sweep is specified, we set *h*_*b*_ = 0. Moderate offsets preserve or strengthen individual bias, whereas sufficiently large offsets suppress responding (Supplementary Note S4).

Except where noted, the connectivity is shaped by experience rather than prescribed. Training and learning are one process here, and we use the two words interchangeably. A network is trained by running it with plasticity switched on, and the learning is unsupervised. The network starts with no recurrent connections, and during training it pursues a single target that performs an uncorrelated random walk across a square arena of side 30 units. The arena is small so that the agent stays near the target and must repeatedly re-orient to it rather than spend the run in transit; it nonetheless touches a wall on under 2% of steps after the transient (Supplementary Note S3). Its four walls are hard boundaries, so any step that would move the agent or target outside the arena is clipped at the boundary in each affected coordinate. The same rule applies in the larger arena used for the choice tasks. Every network starts from the same geometry, as specified in the Methods. Because sensory input is topographic, a target at a given bearing activates the contiguous group of neurons tuned near it, so that as the agent pursues the target, neurons with similar preferred directions are repeatedly in the same state and a Hebbian rule on the neural states strengthens their coupling. Neurons with distant preferred directions are repeatedly in opposite states, and their coupling decreases below zero, becoming inhibitory. Hebbian learning spontaneously builds a center–surround profile resembling the prescribed ring of Eq. 1, together with departures from it that differ from network to network. Pursuit of a single target is the only task an agent is ever trained on. No agent has experienced multiple targets before being tested on them, and every post-training behavioral result outside the continued-plasticity study of Section II I comes from a connectivity frozen once training ends. Two variants extend this protocol, a weak plasticity left on after training ends so that the weights keep changing while the agent behaves, and a training run that begins from a cosine ring rather than from zero. The rule, the arena and the parameters are given in the Methods, *Connectivity and Hebbian learning*.

Alongside these learned networks, we also examine networks with *prescribed* heterogeneity introduced directly into their connectivity rather than generated through learning, which lets us separate effects that require learning from those that follow from heterogeneity however it arises. One construction adds an independent random number, drawn uniformly between − 1 and 1, to every weight of the cosine ring and clips the result to the same range. This changes the strength of each connection and, where the draw is large enough, its sign, so a connection that was excitatory can become inhibitory. The other leaves the ring form intact and gives each neuron its own tuning exponent *ν*_*i*_, a per-neuron spread around *ν* = 0.5, so that neurons differ in how broadly they couple. In comparisons with normalized reference networks, the added noise is instead Gaussian with standard deviation 0.3, and both kinds of prescribed network are rescaled to a common reference gain, while learned networks retain their own gain (Methods, *Connectivity*).

In every network the neurons lie on a ring, and only the connectivity between them varies. A *perfect ring* is one whose weight between two neurons depends only on the angular separation of their preferred directions; its weights differ widely, but every neuron has the same profile. Connectivity is *heterogeneous* when its weights depart from that profile neuron by neuron. We decompose any connectivity as *J* = *C* + *R*, with a network-specific *circulant backbone C* shared across neurons within that network and a *residual R* containing the neuron-specific departures. The backbone is obtained by averaging the weights over all neuron pairs at each angular offset. Each network therefore has its own backbone; these backbones share a center–surround organization but need not have identical weights. The residual has two properties that play different roles here, its *amount*, the overall size of those departures, and its *arrangement*, their specific pattern. Overall coupling strength is the *gain*, the Frobenius norm (the root of the summed squared weights) of *J*; a *gain-matched* control is rescaled to a reference gain so that added structure is not mistaken for stronger coupling.

Each agent is scored on three spatial tasks. *Single-target tracking* asks the agent to pursue one moving target, scored by how closely it follows. The *equal-target choice* places two equally attractive targets symmetrically and scores whether the agent settles on one of them, a test of commitment and individuality. The *unequal-target choice* makes one target more attractive than the other (of higher quality) by raising its stimulus intensity by half; so that a fixed side preference is not mistaken for skill, we score *orientation-balanced accuracy*, the first-entry accuracy averaged over mirrored right-stronger and left-stronger versions of the task. An agent’s *bias* is its side preference, read from the time it spends in each target zone. The *signed* bias gives which side it prefers, the *absolute* bias how strongly. A trait is *repeatable*, and thus a mark of *individuality*, when it recurs across an agent’s own runs while differing between agents, which we quantify by the intraclass correlation ICC [66, 67], the fraction of the total variance that lies between agents rather than within an agent’s own repeats, so that 1 marks a perfectly reproducible trait and 0 no repeatability at all. Every agent is trained from the same starting geometry, which gives the whole population a shared directional bias, a preference for one direction in the world frame.

### B. The homogeneous agent

The homogeneous model provides a reference for the learned networks, with recurrent connectivity fixed to the same symmetric cosine ring in every agent. Agents differ only in their randomly initialized neural states and the stochastic order and outcomes of neural updates. We first characterize spontaneous movement in the absence of targets, then examine single-target pursuit and choices between two stationary targets of equal or unequal attractiveness. From Section II C onward, we evaluate the learned networks on these three tasks to determine how learning-generated heterogeneity changes their behavior.

#### 1. Spontaneous motion and single-target tracking

In the absence of a target, the symmetric cosine ring’s dynamics depend on the inverse noise parameter *β*, which controls how reliably neurons respond to their recurrent inputs. The network passes through an order–disorder transition as *β* grows [59], located near *β* ≈ 2.7 for this symmetric connectivity (Supplementary Note S4). Below the transition, neural activity is disordered and the agent performs a random walk. Well above it, a stable activity bump produces straight trajectories at nearly constant speed, limiting the model’s ability to capture variation in animal movement. Near the transition, neural noise causes the bump to shift around the ring, producing curved trajectories and occasional reversals.

Introducing a moving target elicits directed pursuit, with trajectories becoming straighter and movement faster as *β* increases through the transition. The target takes a step of *v*_*t*_ = 1 in each coordinate, while the agent’s movement scale is *v*_0_ = 10. Its realized speed depends on the neural population vector (Methods). At each step both of the target’s coordinates change by *v*_*t*_, each with an independent random sign, so away from the walls every step is diagonal, of length 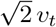. This test differs from training (Section II A), in which the agent also pursues a single moving target, but with plasticity switched on and with each coordinate of the target taking a Gaussian step of standard deviation *v*_*t*_ (Methods). The time-averaged distance to the target is smallest near the transition [59] (Supplementary Note S5), because a weakly ordered network is more responsive to a stimulus and re-orients readily, though it also travels more slowly. The balance between movement speed and responsiveness depends on the target’s motion. Rapid changes in target bearing favor frequent reorientation, whereas a target that remains near its starting position favors a faster, more direct approach.

The dependence of pursuit on neural noise raises the possibility that selection could tune the noise level to the target’s motion. Although *β* is fixed in the main analyses, in an evolutionary model introduced in Supplementary Note S6 it is inherited and subject to mutation, with reproductive success determined by pursuit performance. Over 100 generations, the evolved mean *β* is higher for rapidly moving than for slowly moving targets at both network sizes. The evolutionary model uses a cosine kernel with *ν* = 1, whose transition differs from the *ν* = 0.5 reference above; the finite runs do not establish convergence to criticality.

#### 2. Choosing between two equal targets

We next replace the moving target with two stationary targets positioned 50 units apart in a square arena with sides of 100 units. Any movement beyond the arena is clipped at the boundary in each affected coordinate. The agent starts below the targets on their perpendicular bisector, equidistant from both, and reaches a target when it enters a circle of radius 10 units around it (Methods). When the two targets are equally attractive, the summed proximity to the pair, ⟨1*/d*_1_ + 1*/d*_2_ ⟩, is greatest at an intermediate noise level, whether the stimulus intensity is uniform or decays with the agent’s distance to each target (Supplementary Fig. S2). Proximity to the targets, however, does not imply sustained commitment to either one. The homogeneous agent initially approaches one target through the geometry-dependent transition described in the Introduction. As the agent approaches the pair, their angular separation increases, and the network shifts from a compromise heading between them to a heading toward one target, a transition also observed in animals [55, 58]. This initial selection does not produce sustained commitment, as the agent fails to remain near the target after reaching it. At small *β* the agent moves slowly and irregularly between the targets. At larger *β* it repeatedly crosses from one to the other. Because sensory input encodes target bearing without distance, reaching a target provides no explicit signal to stop, and the agent continues moving (Supplementary Note S5). These behaviors illustrate two forms of failure to maintain a choice between equally attractive alternatives [68]. The repeated switching resembles the back-and-forth walking of flies between inaccessible visual targets in Buridan’s paradigm [69, 70]. This lack of sustained commitment persists across a broad range of inverse noise *β* and input offset *h*_*b*_ (Supplementary Note S4).

#### 3. Choosing between two unequal targets

When one target produces a stronger sensory input than the other, the homogeneous agent reliably reaches the more attractive target first (Supplementary Note S5). It can therefore discriminate between unequal alternatives despite failing to maintain commitment when they are equally attractive. Section II G examines how this accuracy depends on neural noise and compares performance across homogeneous and heterogeneous networks. The homogeneous agent thus performs well in pursuit and unequal-target choice but fails to sustain a choice between equal targets. Its symmetric connectivity provides no persistent internal preference for either alternative, and the identical connectivity across agents cannot support stable individual differences. We next examine how Hebbian learning changes this connectivity and generates variation between agents (Section II C).

### C. Hebbian learning self-organizes structured, heterogeneous connectivity

#### 1. Connectivity built by experience

During training, agents pursue a single moving target while Hebbian learning modifies their recurrent connection weights. In this task, nearby neurons tend to receive similar input and occupy the same state, generating correlations that Hebbian learning uses to modify their recurrent connections (Methods).

All agents begin training at the same position relative to the target, the agent at a corner of the training arena and the target at its center, so that the target initially lies at a bearing of 45^*°*^ and subsequently performs a random walk. This shared starting geometry produces a modest directional bias across the trained population, as examined in Section II H. Individual agents nevertheless develop distinct preferences, and our primary focus is on these differences between agents.

We first examine the connectivity produced by learning and how well the trained agents track a moving target. The standard ensemble’s development is examined in Section II H; Fig. 2A–E characterize separate connectivity-emergence protocols summarized in Supplementary Table S40. We then assess their choices between targets of equal attractiveness in Section II D and unequal attractiveness in Section II F.

**FIG. 2.**
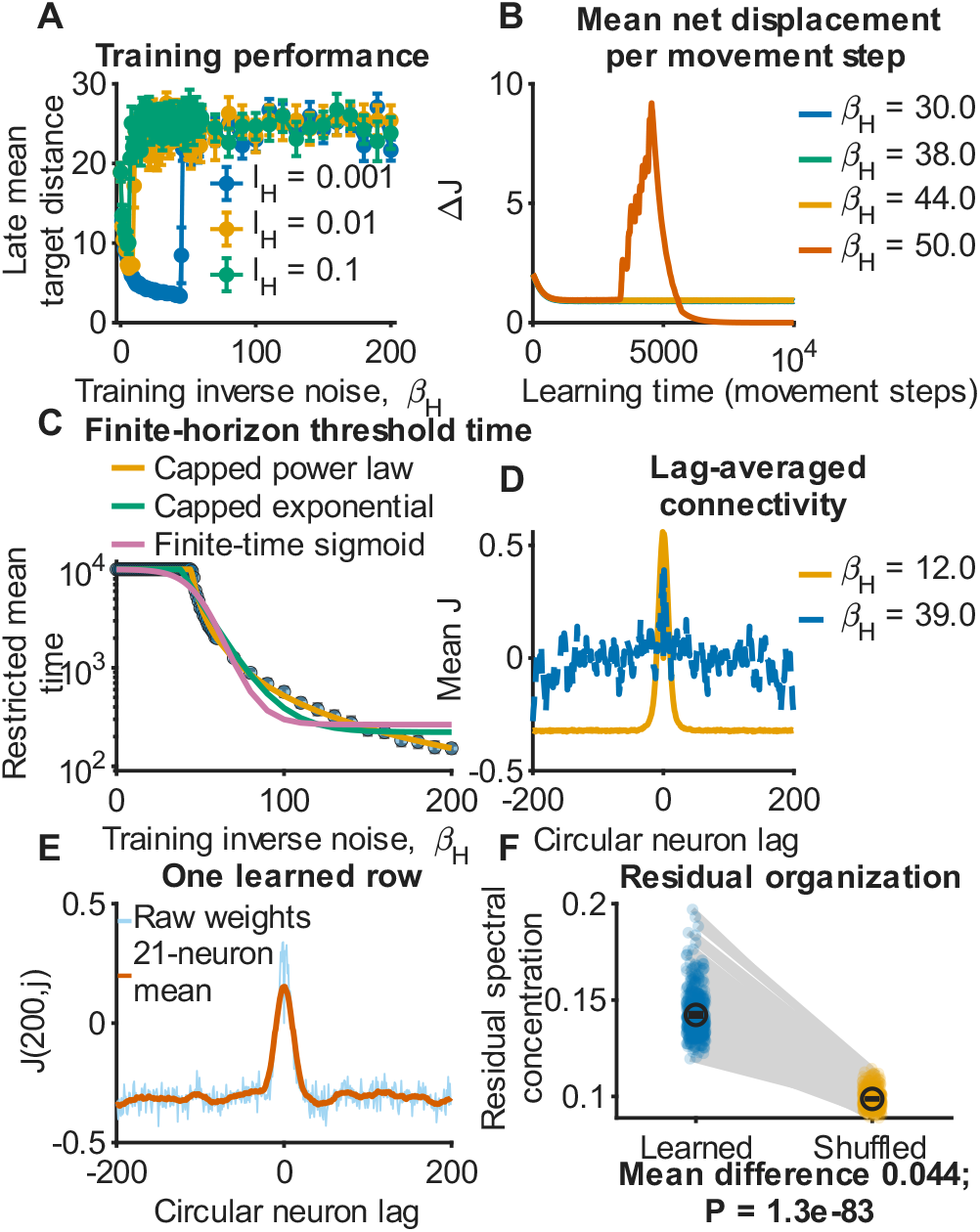
Spin-correlation Hebbian learning self-organizes structured, quasi-periodic connectivity. (A) Late mean distance between agent and target in the learning runs as the training inverse noise *β*_*H*_ is varied, at the indicated learning rates. (B) Net connectivity displacement per movement step, Δ*J* (*t*) = ∑ _*ij*_ |*J*_*ij*_ (*t* + 1) *− J*_*ij*_ (*t*)|, averaged over 10 independently trained networks at each indicated training inverse noise. Mean displacement remains low for an initial period, then rises sharply at *β*_*H*_ = 50; the lower-*β*_*H*_ curves remain low over the plotted horizon. (C) Restricted mean threshold-crossing time under the 10,000-step horizon, with non-crossing runs assigned the horizon. Three horizon-capped fits are drawn, a power law, an exponential and a finite-time sigmoid, showing that the decline does not depend on the form assumed. The fit describes the quasistationary lifetime and does not establish a thermodynamic divergence. (D) Lag-averaged connectivity from one learned network under each of two illustrative *N* = 400 training conditions. The lower-*β*_*H*_ profile is center–surround; the higher-*β*_*H*_ profile is one-dimensional and quasi-periodic. Both noise and sensory width differ between the conditions, so this comparison does not isolate either parameter’s effect. No behavioral result is drawn from the saturated example. Raw and locally smoothed weights in one representative row of a center–surround learned connectivity, illustrating fine-scale variation in a neuron’s incoming weights. (F) Spectral concentration in the residual *R* = *J − C* across 500 learned networks and their exact long-range-shuffle controls. Points are networks; markers and bars show means and 95% network-bootstrap intervals. The shuffle preserves each row’s protected local entries and its long-range weight multiset. Learning uses zero input offset throughout; the offset *h*_*b*_ enters only when a trained network is probed, where it modulates neural input without changing the connectivity norm (Supplementary Fig. S12). Panels A–C use *N* = 100; panels D and E use *N* = 400, so their lag axes run to *±* 200. Panel E and the *β*_*H*_ = 12 profile of D use receptive-field width *σ* = 8*π/N* ; the *β*_*H*_ = 39 profile uses *σ* = 2*π/N*. Panels A–E use connectivity-emergence protocols detailed in the Methods, with learning and integration settings differing from standard training. The threshold 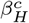 is fitted only to the protocol of A–C and does not apply to the illustrative conditions in D and E. Panel F uses the standard training task.

#### 2. A threshold in the training noise

Neural noise during training determines whether learning produces connectivity that supports effective pursuit. We quantify tracking performance by the mean distance between agent and target during the final portion of training, with smaller distances indicating closer tracking. As the training inverse noise *β*_*H*_ increases, neural noise decreases and tracking initially improves. Beyond a threshold, however, the mean distance rises sharply and tracking performance deteriorates (Fig. 2A).

Changes in connectivity during training reveal the structural reorganization associated with this loss of tracking performance. After an initial adjustment, the weights remain relatively stable for an extended period before undergoing rapid change (Fig. 2B). This reorganization arises from the Hebbian response to correlations between neural states, without any reward or feedback on pursuit performance. The duration of the relatively stable period decreases sharply as *β*_*H*_ increases beyond a threshold, which we denote 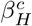 (Fig. 2C). We estimate this duration using fits capped at the maximum training time, assigning that limit to runs in which no reorganization occurs. The resulting timescale describes the persistence of the connectivity within the observation period and does not establish a thermodynamic divergence.

The training runs in Fig. 2A–C differ from the standard protocol in their learning and forgetting rates, learning-time inhibition, and number of neural updates per movement step. The fitted threshold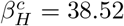 therefore applies specifically to those conditions. Under the standard protocol, tracking deteriorates at substantially lower *β*_*H*_. Mean post-training tracking error increases from 3.5 at *β*_*H*_ = 14 to 12.4 at the standard value of *β*_*H*_ = 17, and reaches 37.9 at *β*_*H*_ = 21, where pursuit is severely impaired. The networks used in the subsequent analyses are thus trained in a regime where tracking has already deteriorated, although it has not reached the severe impairment observed at higher *β*_*H*_ (Supplementary Note S7).

#### 3. Learning produces either center–surround or saturated connectivity

Learning can produce local excitation and longer-range inhibition. Neurons with nearby preferred directions become strongly and positively connected, and the coupling weakens and then becomes inhibitory as their preferred directions differ more. This is the center–surround profile commonly assumed for ring-attractor networks [20, 53] (Fig. 2D); Hebbian learning has been shown to produce it in head-direction models [34]. Under other training conditions, continued learning drives the weights to their imposed bounds, producing a saturated network whose mean connectivity varies quasi-periodically with angular separation (Fig. 2D). These two illustrative *N* = 400 networks differ in both training noise and sensory width, so they do not isolate either parameter’s effect or establish a common threshold. Saturation is accompanied by a loss of effective tracking. We therefore focus the subsequent behavioral analyses on networks that develop center–surround connectivity, except where we explicitly examine saturation and its consequences.

Periodic connectivity of this kind has previously been obtained by training networks toward a spatial objective [27–29]. Here no objective or error signal is supplied, and the periodicity self-organizes under an unsupervised rule during pursuit training, as periodic structure has in earlier work under Hebbian learning with adaptation [36] and under spike-timing-dependent plasticity in an initially unstructured recurrent network [35]. We describe it as one-dimensional quasi-periodic connectivity and draw no inference about grid cells, since the model represents a single angular variable, produces neither two-dimensional rate maps nor grid scores, and does not navigate in the regime where the periodicity appears.

#### 4. A structured residual unique to each agent

Averaging connectivity across neurons obscures differences in their individual connection patterns. In networks with a center–surround profile, individual neurons’ incoming weights show structured deviations from that network’s circulant backbone defined in Section II A. Figure 2E illustrates this fine structure by comparing one neuron’s incoming weights with a local moving average along the ring, using the window width indicated in the panel. Across the neural population, these deviations show structured angular variation beyond the network’s mean center–surround profile.

Across the 500 learned networks, the center–surround backbone accounts for approximately 91% of the total squared connection weight, measured by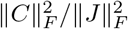.

We construct this backbone by averaging weights between neuron pairs with the same directed separation around the ring. Writing this separation as *d* = (*j*−*i*) mod *N*, its entries are 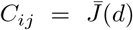, where 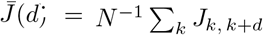 and indices are taken modulo *N*. The residual *R* = *J* − *C* accounts for the remaining 9%. Although small in amplitude, with a median within-network standard deviation of 0.0573, it contains structured differences between neurons.

Self-connections are excluded by setting *J*_*ii*_ = 0. Reciprocal connections generally differ, with a median normalized asymmetry ∥*J*− *J*^T^∥ _*F*_ / ∥*J* ∥_*F*_ of 0.529 across networks (interquartile range 0.5251–0.5341). This corresponds to approximately 7% of the total squared connection weight in the antisymmetric component of *J*. The asymmetry arises because each Hebbian update modifies the incoming weights of a single postsynaptic neuron. Reciprocal weights are therefore updated at different times and can reflect different neural states. Such asymmetry is biologically plausible because reciprocal synapses are distinct connections, and mechanisms such as spike-timingdependent plasticity can also produce unequal reciprocal weights [71, 72].

The residual exhibits quasi-periodic variation around the ring, with Fourier power concentrated at particular angular frequencies. We quantify this structure using spectral concentration, defined for each neuron as the fraction of residual power at its strongest frequency and then averaged across neurons within each network. Mean spectral concentration is 0.142 in learned networks and 0.099 in matched controls that shuffle each neuron’s longrange incoming weights while preserving their values and leaving local connections unchanged (Fig. 2F). The mean paired difference is 0.0436 (95% interval 0.0429–0.0444), a 44% increase over the control. Spectral concentration is higher in every one of the 500 learned networks than in its matched control (signed-rank *P <* 10^*−*80^), demonstrating a consistent, though modest in absolute magnitude, departure from shuffled connectivity. Smoothing the residual increases spectral concentration to 0.295, providing a comparison in which broad angular structure is retained and finer fluctuations are suppressed. The learned residual therefore has greater spectral concentration than the shuffled control but substantially less than its smoothed counterpart (Supplementary Note S7). Because shuffling also removes smooth autocorrelation, this comparison does not uniquely identify periodic organization; quasi-periodicity is a descriptive characterization, not an isolated cause of the behavioral effects.

Neurons differ from one another in this respect. Some show a clear quasi-periodic modulation and others do not. Quasi-periodic modulation was more prevalent in the larger network: Fisher’s *g* exceeded the significance threshold in 32% of neurons at *N* = 100, averaged across 500 networks, and in 51% in the *N* = 400 example. Training parameters and the exclusion scale also differed, so this comparison does not isolate the effect of network size. The matched long-range reshuffles give 5% and 4%, respectively. This excess spectral concentration is present in a minority of neurons at *N* = 100 (Supplementary Note S7 and Supplementary Fig. S3). This fine modulation, several alternating positive and negative lobes around the ring rather than a single bump, is superimposed on the broad center–surround profile and resembles the detailed connectivity of biological navigation circuits, whose fine organization is thought to help a single population represent more than one angular variable at once, although its computational role is still only partly understood [73]. Two further features of learned networks are reported in the Supplementary Information, namely multiple peaks in neural activity across movement directions, and a tendency of a minority of agents to return repeatedly to the same spot away from either target (Supplementary Note S8 and Supplementary Fig. S4).

Hebbian learning thus produces structured variation between neurons and between agents that is absent from the homogeneous ring. We next examine how this variation relates to repeatable preferences between equal targets (Section II D). We then manipulate the learned connectivity to distinguish the behavioral effects of the amount of heterogeneity from those of its spatial arrangement (Section II E).

### D. Learned networks make repeatable, individual-specific choices between equal targets

#### 1. Learning leads to idiosyncratic behavior

We next test whether agents trained solely on singletarget pursuit (Section II C) develop consistent preferences between two equally attractive targets. Unlike homogeneous agents, which can reach a target, but not remain there, and exhibit no side preference (Section II B), learned agents show both sustained commitment and repeatable preferences. Their neural activity remains localized but fluctuates irregularly, supporting persistent movement toward a preferred target across independent runs (Supplementary Fig. S5). Pooled over 100 trained agents with five independent test runs each, occupancy concentrates at the target zones (Fig. 3A,B).

**FIG. 3.**
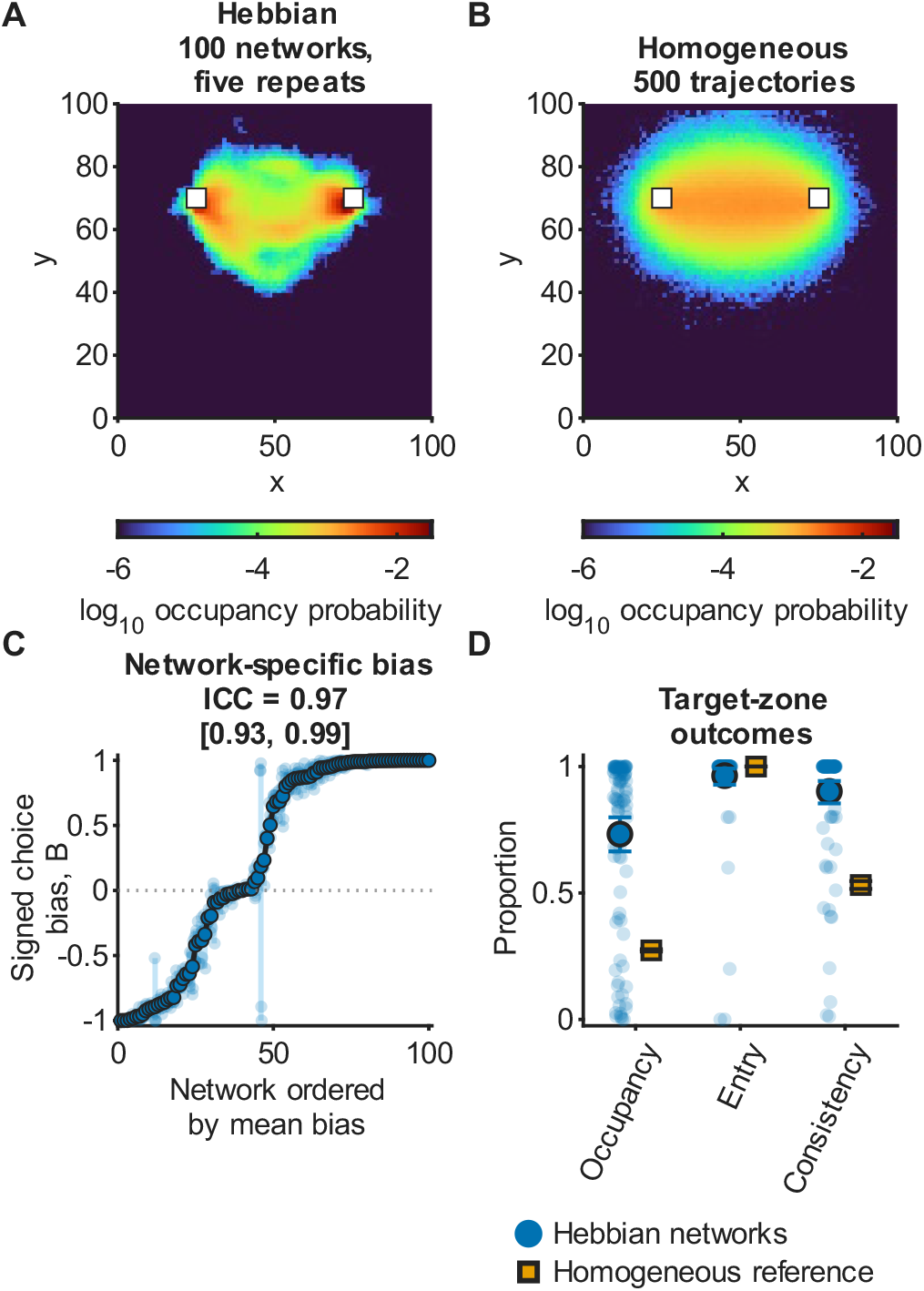
Learned networks make repeatable, individualspecific choices between equal targets. (A) Pooled occupancy in the equal two-target choice for 100 learned networks with five repeats each. Occupancy concentrates at the two target zones. White squares in A and B mark the two equal targets. (B) Pooled occupancy for 500 independently seeded trajectories of the homogeneous *ν* = 0.5 reference, each network type at the inverse noise at which it is reported, *β* = 40 for the learned networks in A and *β* = 4 for the homogeneous ring in B. The homogeneous agent approaches the pair but settles on neither, and its occupancy stays in the region between the targets. Panels A and B share logarithmic color limits. Trajectories of representative learned networks are shown in Supplementary Fig. S5. (C) Signed occupancy bias *B* = *O*_*R*_ *− O*_*L*_, the post-transient difference between right- and lefttarget-zone occupancy, for five repeats of each learned network, ordered by network mean bias. Vertical ranges and faint points show repeats; the dark line shows network means. ICC and its 95% network-bootstrap interval are printed in the panel. (D) Posttransient target-zone occupancy, complete-trajectory target entry, and consistency among target-zone visits for learned and homogeneous networks. Markers and bars show means and 95% bootstrap intervals; faint points show learned-network means.

The side an agent chooses is a stable property of that agent. Repeated tests of the same agent return to the same target, while different agents can prefer opposite sides. The individual biases span the whole range from a strong left to a strong right preference and are highly reproducible, with a spread between agents far exceeding the variability within an agent. At *β* = 40 the signed occupancy bias, the post-transient difference between time spent in the right and left target zones (Supplementary Note S9), is highly repeatable, with ICC = 0.974 and a 95% interval of 0.932–0.995 (Fig. 3C), and 0.975 at *β* = 60; the full ensemble also spans a broad distribution of preferences (Supplementary Fig. S6).

The proportion of trajectories entering either target zone is similar for learned and homogeneous networks, but learned agents spend more time near the targets and return more consistently to the one first selected (Fig. 3D). At *β* = 40 (Methods, Table I), learned agents spend an average of 73.3% of the post-transient period within the target zones, and only 3.6% of trajectories reach neither zone. Learning thus produces connectivity that supports sustained, individual-specific preferences in a choice task the agents have not previously encountered. Repeatability holds over a broad range of the inverse noise rather than at the single value tested above. It rises from near zero in the low-*β* regime, where commitment is unstable, through 0.82 at *β* = 16 and 0.92 at *β* = 20, and plateaus above 0.9 from *β* = 40 onward (Supplementary Fig. S7). Because the preferred side differs from agent to agent, a pooled occupancy map alone understates how strong and how repeatable each agent’s own preference is.

**TABLE I.** Selected fixed-*β* comparisons and dynamical peak estimates for the four network types. Noise sweeps and the separate gain-matched Gaussian-noise controls are specified in Supplementary Note S4. Susceptibility and amplitude-relaxation peaks are stimulus-free diagnostics. Only the symmetric ring has an equilibrium transition, and its susceptibility and relaxation peaks lie close together; for the other three they are reported separately and neither is treated as locating a critical point. Learned and clipped-uniform values use network means over three repeats on the complete 49-point grid ending at *β* = 200; 5*/*20 learned and 4*/*20 clipped-uniform relaxation maxima remain at its upper boundary. The values in the second column are not equally far from where each type’s diagnostics peak, so magnitudes should not be compared across types.

| Network type | Selected fixed-β comparisons | Susceptibility peak | Relaxation peak |
| --- | --- | --- | --- |
| Symmetric cosine ring, $\nu = 0.5$ (equilibrium transition $\beta \approx 2.7$) | sweeps; 40 and 60 for matched controls | 2.5–3.0 | 3.0–3.5 |
| Learned Hebbian | 40 and 60 | median 58 (IQR 43–75) | median 120 (IQR 100–175) |
| Clipped uniform-noise | 16 | median 3.5 | median 44 (IQR 13.25–110) |
| Heterogeneous tuning | 4 | not separately estimated | not separately estimated |

The repeatable differences between agents are encoded in their learned connectivity, consistent with developmental stochasticity producing stable behavioral idiosyncrasies [6, 40].

The individual differences extend beyond target choice to how agents move. Moving without a target, learned agents differ in mean speed, in how widely their step speeds are spread about that mean, measured as the coefficient of variation, and in how sharply they turn, measured as the mean absolute change of heading per step. Each of these three descriptors is almost perfectly reproducible within an agent across independent runs, with an intraclass correlation ICC ≥ 0.99 for all three and a spread between agents far exceeding the within-agent variability, whereas the homogeneous ring moves in much the same way on every run (Supplementary Fig. S8). Learning thus gives each agent an individual movement style, of which the target preference is one part. In this sense, individuality is already present in free behavior, so what the training task leaves open is specifically the left–right side preference and not individuality in general (Section II H).

#### 2. Heterogeneous weights produce individuality

Heterogeneity in the weights, which learning generates naturally, underlies the observed biases. To ask whether this heterogeneity must come from learning, we tested agents that never learned but carry prescribed heterogeneity instead. The tested heterogeneity constructions produce side preferences, whether acquired by learning, introduced as random noise on a cosine ring, or generated by heterogeneous directional tuning. Repeatable preferences therefore arise under several tested forms of heterogeneity and do not require learning (Supplementary Note S5). With their coupling gain normalized to the common reference, these prescribed agents also show high across-run repeatability, with ICCs of 0.82 and 0.95 at *β* = 16 and approximately 0.89–0.97 over *β* = 20–60 (Supplementary Fig. S7).

The random-noise agents used as the secondary comparison in Section II F were left at their own coupling strength instead of being normalized to the common reference gain, and at that strength they commit only weakly. At *β* = 16 their absolute bias stays below 0.06, their target occupancy is 0.067, they fail to decide on of runs, and their repeatability is only ICC = 0.414. They are included because they carry a directional preference without any learned structure, which shows that the side advantage of Section II F follows from having a preference at all rather than from having learned one. These agents use the clipped-uniform construction, which adds uniform noise on [− 1, 1] to each weight of the ring and leaves the coupling scale free (Methods). The matched-gain random-noise agents quoted above use a different construction, Gaussian noise of standard deviation 0.3 with the whole matrix then rescaled to the Frobenius norm of 19.6 (Supplementary Fig. S7).

Learning is therefore not the only route to a reliably biased agent, since prescribed heterogeneity reaches the same repeatability. What learning adds is that the preference is acquired from the agent’s own experience rather than engineered by hand. Section II E tests each modified connectivity on all three tasks to ask whether the particular pattern of learned weights matters, or only the degree of heterogeneity. The third task, choosing between unequal targets, also addresses a separate question— whether this heterogeneity helps the agent choose the stronger of two (Section II F).

### E. Rearranging learned connectivity retains repeatable preferences and improves pursuit

When two targets are equally attractive, neither target is objectively preferable; the equal-choice task tests whether the agent sustains commitment to either target. The homogeneous ring can reach a target but cannot remain near it (Section II B), whereas a learned network can (Section II D). This raises two questions. Does remaining near the target depend on the specific pattern of weights produced by learning, or would weight variation of the same magnitude suffice? And does retaining that pattern come at a cost in the task the agent was trained to perform, following a single moving target?

Comparing a learned network with a homogeneous ring answers neither question, because the comparison changes the backbone, the variation of individual weights around it, and the overall coupling strength at the same time. We therefore altered the weights of each learned network while holding the other properties fixed (Fig. 4A,B). We decomposed each connectivity as *J* = *C* + *R*, where *C* is that network’s circulant backbone and *R* is its residual (Section II A), then rebuilt the network from its own *C* and a rearranged *R* in three ways. In the first, at each angular offset, we assigned the same residual values to different pairs of neurons. In the second we kept the size of each wave in the residual pattern but shifted its peaks around the ring. In the third we replaced the learned residual with a random residual of the same size, either completely or as the intermediate mixture (*λ* = 0.5) of the two (Methods, Eq. 12). Each rearrangement preserved the mean and spread of the residuals at every offset and the network’s total weight strength, changing only how the residuals were arranged (Supplementary Note S10). We then tested each rebuilt network independently on pursuit of a single moving target, equal choice, and unequal choice, using 100 learned networks and three runs of each at inverse noise *β* = 60. When given two equally attractive targets, agents with rearranged connectivity retain substantial preferences and high repeatability. Preference strength is 0.554 when none of the learned pattern remains and 0.515 for the intermediate mixture (*λ* = 0.5), compared with 0.537 for the original learned networks; these bias magnitudes are shown in Fig. 4C. Side preference remains highly repeatable after rearrangement, with ICCs of 0.987 for the reshuffled version, 0.986 for the wave-shifted version, and 0.963 for the intermediate mixture (*λ* = 0.5), compared with 0.978 for the learned networks (Supplementary Note S10). Removing the deviations altogether, leaving only the backbone, nearly eliminates the preference. Its strength falls to 0.055 and its repeatability to 0.029. Restoring the backbone to the network’s original total weight does not recover the preference; learned networks still exceed this rescaled control by 0.485 (95% interval 0.412–0.557). The learned arrangement also takes longer to reach either target for the first time, 153.54 movement steps compared with 66.97–70.44 after rearrangement. Paired differences in preference strength were 0.017 [−0.108, 0.074] for learned minus moment-matched control (*λ* = 0), −0.038 [−0.130, 0.054] for learned minus shuffled residual, and− 0.073 [−0.164, 0.017] for learned minus phase-randomized residual. Thus, the learned pattern is not required for commitment, although these intervals leave room for differences in its strength.

**FIG. 4.**
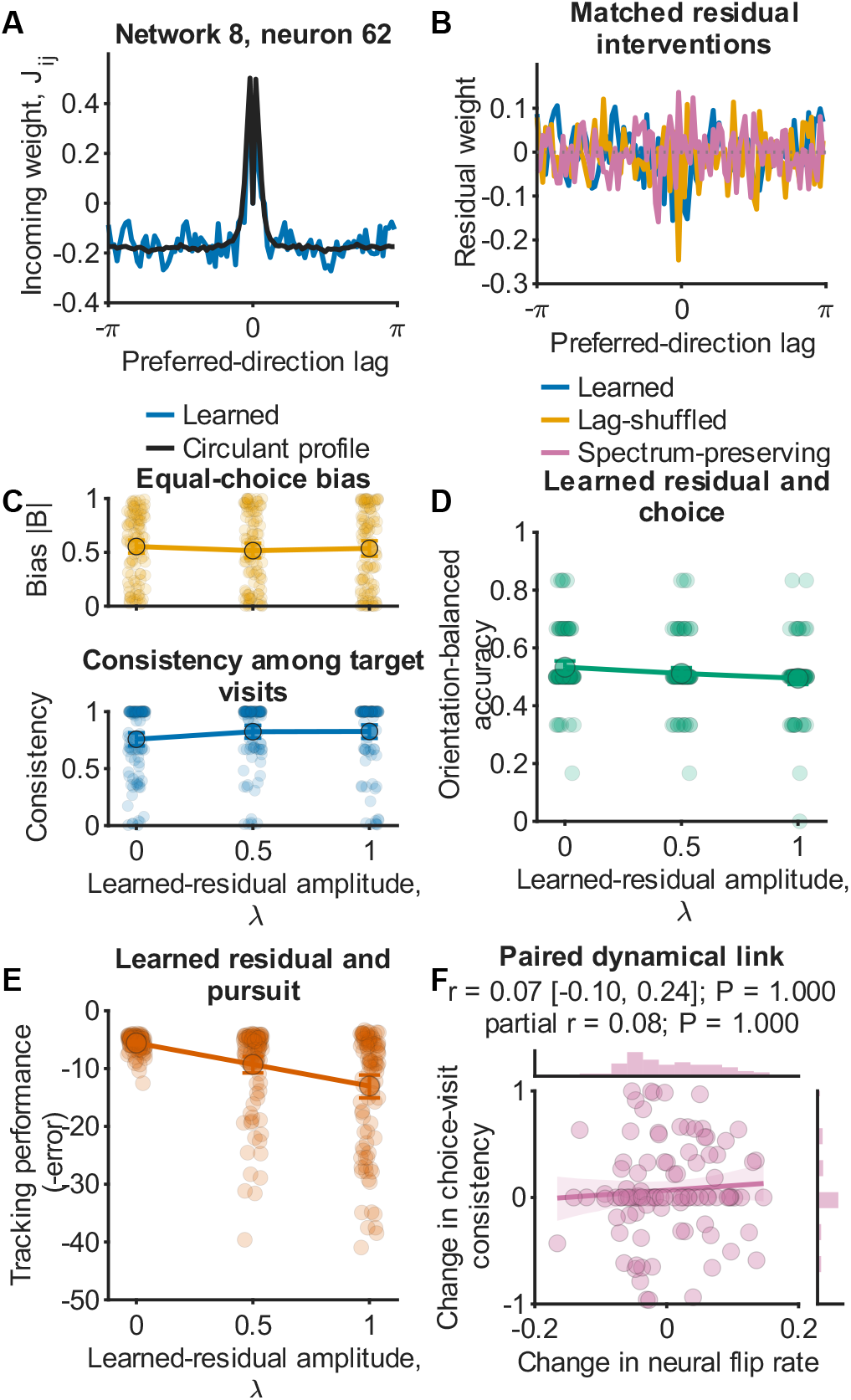
Residual heterogeneity supports repeatable preferences; its arrangement determines pursuit performance. (A) Incoming weights for a representative neuron in one learned connectivity and its directed-lag circulant profile. (B) The corresponding learned residual in blue, a shuffle that assigns each lag’s values to different pairs of neurons in orange, and a phase control in pink that keeps the amplitude of each lag’s waves but shifts their peaks around the ring. (C) Absolute equal-choice bias (upper) and consistency among target-zone visits (lower) as the learned residual is replaced by random values of the same size, 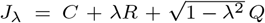. At the intermediate mixture (*λ* = 0.5) the learned residual has half its original amplitude and contributes one-quarter of the residual squared norm. (D) Orientation-balanced unequal-choice accuracy and (E) pursuit performance along the same replacement; lower pursuit performance means larger tracking error. (F) Network-level paired change in neural flip rate and choice-visit consistency for learned networks relative to the *λ* = 0 fully replaced counterpart. No association between the two changes was detected across networks: *r* = 0.07 (95% interval *−* 0.10–0.24; Holm-adjusted *P* = 1), and *r* = 0.08 after adjusting for changes in equal-choice mean activity and speed (Holm-adjusted *P* = 1). The panel shows the marginal distributions of the two changes, a least-squares line with its shaded 95% band, and the Pearson estimate with its 95% network-bootstrap interval. Faint points are network means; lines, opaque markers, and bars show population means and 95% network-bootstrap intervals.

When one target is more attractive than the other, we detected no difference in orientation-balanced accuracy between learned and rearranged networks (Fig. 4D; Supplementary Note S10). Nor is poor pursuit a necessary cost of commitment. In a separate training comparison, networks trained at *β*_*H*_ = 14 retain a substantial side preference (0.676, compared with 0.743 at *β*_*H*_ = 17) while tracking a moving target more closely (mean distance 3.55 versus 12.45). These results come from ten seeds per condition and three probe runs per seed (Supplementary Note S7; Supplementary Table S16). Because this comparison uses a different pursuit probe, its tracking distances should be compared with each other, not with those reported above. Learned networks can therefore combine a stable choice with good pursuit.

In pursuit, agents with learned connectivity remain a mean distance of 13.02 from the moving target, compared with 5.57–5.83 for the three rearranged versions— the largest and most consistent difference across the tasks (Fig. 4E; Supplementary Note S10). Neural switching during pursuit was descriptively lower in the learned networks, but the paired intervals included zero, so these measurements do not establish slower switching as the cause. Because the rearranged networks retain the same amount of departure from the backbone yet track about as well as the backbone alone (mean distance 5.34), the poor tracking depends on the learned arrangement of those departures. Variation can still impair pursuit. Adding random variation to a plain cosine ring also worsened mean tracking in the noise-amplitude sweep.

Maintaining a choice and following a moving target place different demands on a network. In paired comparisons of each learned network with its fully rearranged counterpart, the change in neural switching during equal choice shows little association with the change in how consistently the agent returns to its first-chosen target (*r* = 0.07, 95% interval − 0.10–0.24; Fig. 4F). Which behavior matters more will depend on the setting, including the cost of reversing a choice and the need to keep up with a moving target.

### F. Side preference predicts how accuracy depends on target position

#### 1. Heterogeneity narrows the conditions under which the choice stays accurate

Agents repeatedly favor the same side when choosing between two targets (Fig. 3). In animals, consistent behavioral biases have been linked to problem-solving performance [74, 75], raising the question of whether a stronger side preference makes an agent better at decision-making. The answer depends on how accuracy is measured. If the stronger target is always on the right, a right-biased agent may score well even if it ignores target quality. In the primary repeated-trial cohort, we therefore tested each agent in both orientations, placing the stronger target on the right and then on the left, using the same random numbers for both runs. We average accuracy across the two. This orientation-balanced accuracy is the expected score when the better option is equally likely to be on either side, and it is the accuracy we report from here on (Supplementary Note S9). We also distinguish the side an agent prefers from the strength of that preference.

To see how accuracy changes as the choice becomes harder, we varied the target-intensity ratio from 3:1 to nearly even at 1.1:1. At every noise level tested, the homogeneous ring approaches ceiling accuracy as the difference between targets grows. The two prescribed heterogeneous networks improve more slowly, while the learned networks are generally least accurate across the tested ratios, improving only modestly at *β* = 40 and barely at all at *β* = 60 (Supplementary Fig. S9). In this allocentric difficulty comparison, none of the forms of heterogeneity improved average accuracy over the homogeneous ring. They produced repeatable differences between agents but narrowed the conditions under which agents reliably chose the stronger target, with the largest reduction in the learned networks.

#### 2. Side preference predicts how target position affects accuracy

Agents reach the stronger target more often when it is on their preferred side. Among learned agents at *β* = 60, side preference measured when the two targets were equally attractive predicted the difference between accuracy with the stronger target on the right and accuracy with it on the left (*r* = 0.563, 95% interval 0.405– 0.699; Fig. 5A). Agents with prescribed random variation in their weights showed a similar relationship at *β* = 16 (*r* = 0.325, 95% interval 0.116–0.492; Fig. 5B). The learned population favors the right, consistent with the shared initial target bearing during training (Section II H). It reached the stronger target on 0.640 of trials when that target was on the right and 0.362 when it was on the left, giving an orientation-balanced accuracy of 0.501 (Fig. 5C). The position advantage therefore does not depend on learning, although learned agents retain their preferred side much more consistently across runs (Fig. 5D; Section II D). Most agents kept their preferred side as stimulus strength increased, although a minority switched sides as the noise level changed (Supplementary Note S5; Supplementary Fig. S10).

**FIG. 5.**
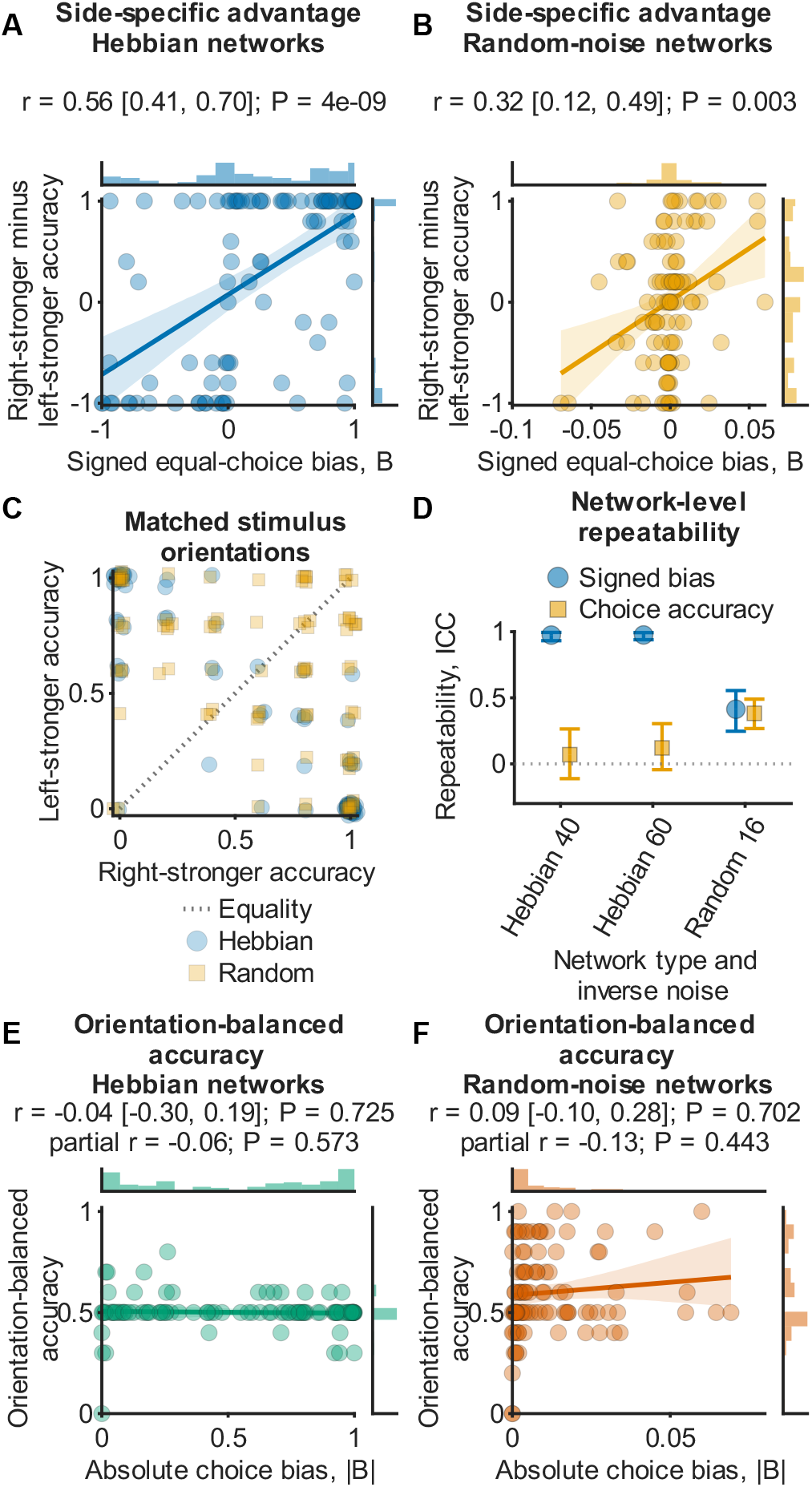
Signed bias predicts the advantage of target position, while preference magnitude shows no detected accuracy association in these allocentric cohorts. (A,B) Signed equal-choice bias versus the difference between right-stronger and left-stronger first-choice accuracy, for Hebbian networks at *β* = 60 (A) and prescribed random-noise networks at *β* = 16 (B). (C) Right-stronger versus left-stronger accuracy; the dotted line is equality. (D) ICC for signed bias and orientation-balanced accuracy in Hebbian *β* = 40, Hebbian *β* = 60, and random-noise *β* = 16 networks. (E,F) Absolute equal-choice bias versus accuracy averaged over the two stronger-target orientations, for the same two network types; no association was detected. Marginal distributions, least-squares lines, Pearson estimates, and 95% networkbootstrap intervals are shown; faint filled markers denote network means. The partial-correlation estimates printed in E and F give the partial *r* of absolute bias with balanced accuracy after adjusting for three-task speed variation, residual variation and connectivity asymmetry. Occupancy-conditional target-choice exclusivity is treated separately in the Supplementary Information. The null in E,F does not generalize across noise levels: a separate allocentric cohort has negative bias–accuracy associations at *β* = 16, 20 and 40 (Supplementary Table S41). The contrasting egocentric results are reported in Section II H 3 and Supplementary Fig. S1.

#### 3. The association between preference strength and accuracy depends on neural noise

We measured the strength of a preference as 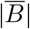, the absolute difference between the fraction of time an agent spends in the right target zone and the fraction it spends in the left. No robust association with orientation-balanced accuracy was detected in the 100-network learned cohort shown in Fig. 5 at *β* = 60 (*r* =− 0.036, 95% interval − 0.298–0.192) and in the randomnoise agents (*r* = 0.094, − 0.097–0.283; Fig. 5E,F). The partial-correlation estimates printed in those panels show no detected association after accounting for speed variation, residual variation and connectivity asymmetry. In the learned cohort, accuracy is close to chance and less repeatable than signed bias; the intervals therefore do not exclude appreciable associations or establish independence at other noise levels.

For the comparison with egocentric networks, we trained 40 allocentric networks under the same standard protocol and tested them across five noise levels. These networks likewise showed no resolved association at *β* = 60 (*r* = 0.094, 95% interval − 0.405 to 0.425). At lower *β*, however, preference magnitude and balanced accuracy were negatively associated: network-level correlations were *r* = − 0.694 (95% interval − 0.841 to −0.513) at *β* = 16 and *r* = − 0.840 (− 0.924 to − 0.730) at *β* = 20. The association was also negative at *β* = 40 (Supplementary Table S41). Thus the high-*β* null does not generalize across operating conditions; these additional correlations are descriptive and use six unequal-choice trials per network.

Chance here depends on how non-entry is treated. Given the observed entry rates, an agent choosing a side at random would score 0.4845 at *β* = 60 and 0.4815 at *β* = 40 (Supplementary Note S11). Learned agents score 0.501 and 0.519, respectively. Neither differs from nominal 0.5 after multiplicity adjustment, although both show a small excess over the entry-aware benchmark in paired comparisons without adjustment for multiple testing. Because network types were tested at different noise levels, their curves in Fig. 6 and Supplementary Fig. S11 provide a more appropriate comparison than isolated scores.

**FIG. 6.**
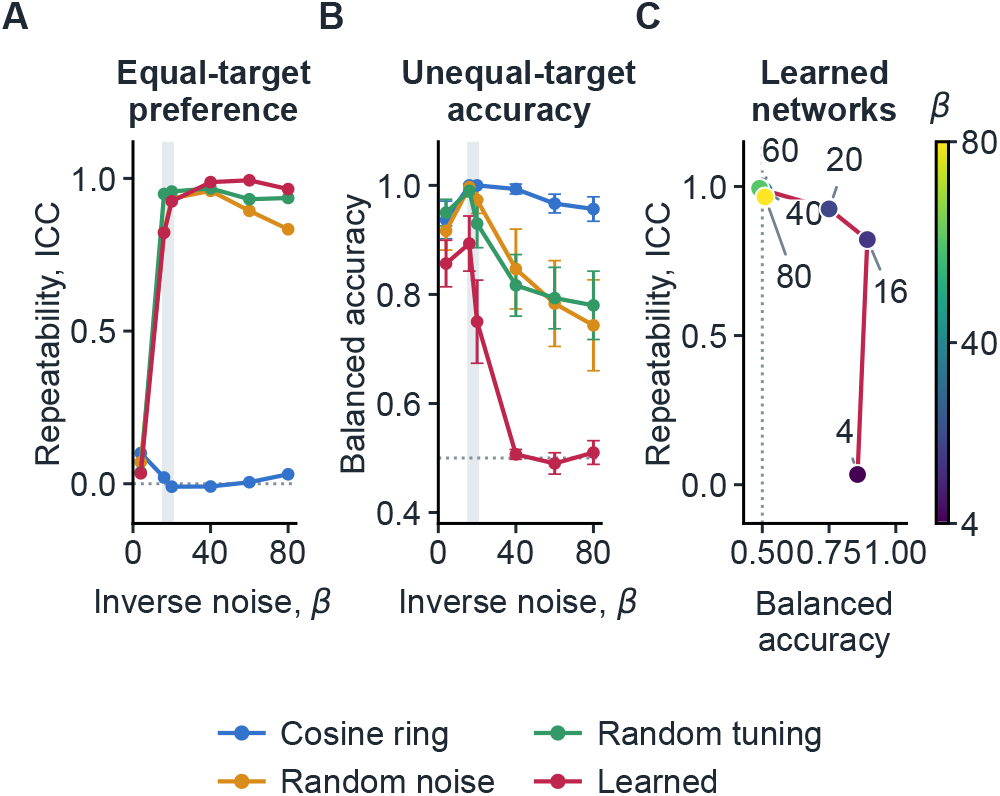
A middle range of neural noise in which learned agents show both repeatable side preferences and accurate choices. All panels use the standard sensory width and zero input offset, the same networks and the same *β* grid, with prescribed types at the fixed Frobenius reference of 19.6 and learned weights unscaled: the homogeneous *ν* = 0.5 cosine ring, prescribed randomnoise and random-tuning networks, and the learned networks. (A) Repeatability ICC of the signed equal-choice bias across five independent runs of each network. (B) Orientation-balanced accuracy on the 1.5:1 unequal choice. Markers and bars are means and*±* 1.96 network standard errors over 30 networks of each type; the dotted line is nominal chance. (C) The two outcomes plotted against each other for the learned networks, with *β* increasing along the curve; marker color gives *β* as in the colorbar, and the dotted vertical line is chance, 0.5. Accuracy and repeatability rise together up to *β* = 16; beyond it accuracy falls as repeatability rises further and then remains near 0.96–0.99, and the shaded band in A and B marks *β ≈* 16–20, where both are high. The cosine ring remains accurate across the tested noise levels, while its side-preference repeatability remains near zero. The inverse noises used elsewhere, *β* = 40 for equal-choice individuality and *β* = 60 for the connectivity interventions, lie to the right of this band.

Occupancy-normalized measures ask how exclusively an agent uses one target among its target visits. In the primary repeated-trial cohort, these measures do not provide a robust relation with balanced accuracy: bootstrap intervals span zero, rank associations are weak, and median held-out prediction does not improve (Supplementary Note S11).

### G. Accuracy and individuality coexist at intermediate neural noise

We varied neural noise in the same learned networks by changing *β*, with higher values corresponding to lower noise. Repeatability of side preference rises with *β* and levels off above 0.9 (Section II D; Fig. 6A), while orientation-balanced accuracy on the 1.5:1 choice declines. Across 30 networks per setting, accuracy falls from 0.89 at *β* = 16 to 0.75 at 20, 0.51 at 40, and near chance at 60. Around *β*≈ 16–20, repeatable preferences coexist with accurate choices (Fig. 6A–C). These population summaries do not imply that every agent in this range has a strong preference.

The homogeneous ring maintains accuracy of at least 0.94 from *β* = 4 to 80, but shows almost no repeatable side preference. Its ICC stays close to zero, ranging from – 0.010 to 0.101. Accuracy declines more gradually in the two prescribed heterogeneous network types, which still score 0.78 and 0.79 at *β* = 60, where learned networks are near chance. Accuracy therefore depends on both connectivity and neural noise.

The analyses of individuality in Fig. 3 and connectivity rearrangements in Fig. 4 use *β* = 40 and 60, respectively. At these settings, side preferences are highly repeatable and spontaneous directional fluctuations are near their peak (Supplementary Note S4; Supplementary Fig. S12). Neither setting gives the highest choice accuracy.

Sensory input width also affects accuracy. The standard field has a standard deviation *σ* = 2*π/N* of one preferred-direction spacing, concentrating input on a few nearby neurons. In a separate sample of 60 learned agents tested in both target orientations at *β* = 60, widening the field to two, four, and eight times its standard width raised balanced accuracy, averaged across the tested intensities, from 0.51 to 0.53, 0.58, and 0.78. The baseline interval, 0.49–0.53, includes chance; the interval at eight times the width is 0.74–0.83. Intensity also affects accuracy, most clearly at the widest field (Supplementary Table S26).

Choice accuracy also depended on the overall neural input. A uniform inhibitory offset could improve accuracy when targets differed in value, although stronger inhibition eventually suppressed responding (Supplementary Note S4; Supplementary Fig. S12). Reduced accuracy therefore depended on the operating conditions and was not an unavoidable consequence of learned individuality.

### H. Training experience shapes individual preferences

#### 1. Individual preferences strengthen with training

During training, directional coherence increased and agents tracked the moving target more closely, while mean angular error decreased modestly from 69^*°*^ to 66^*°*^ (Fig. 7A). Networks trained for longer and then tested with fixed weights showed stronger side preferences, greater variation in signed bias across networks, and fewer trials in which neither target was reached (Fig. 7B). Training thus improved pursuit while generating individual preferences in a separate choice task. Neither the direction of heading bias nor the sign of tracking error during training predicted these later preferences (Supplementary Fig. S13A; Supplementary Note S7).

**FIG. 7.**
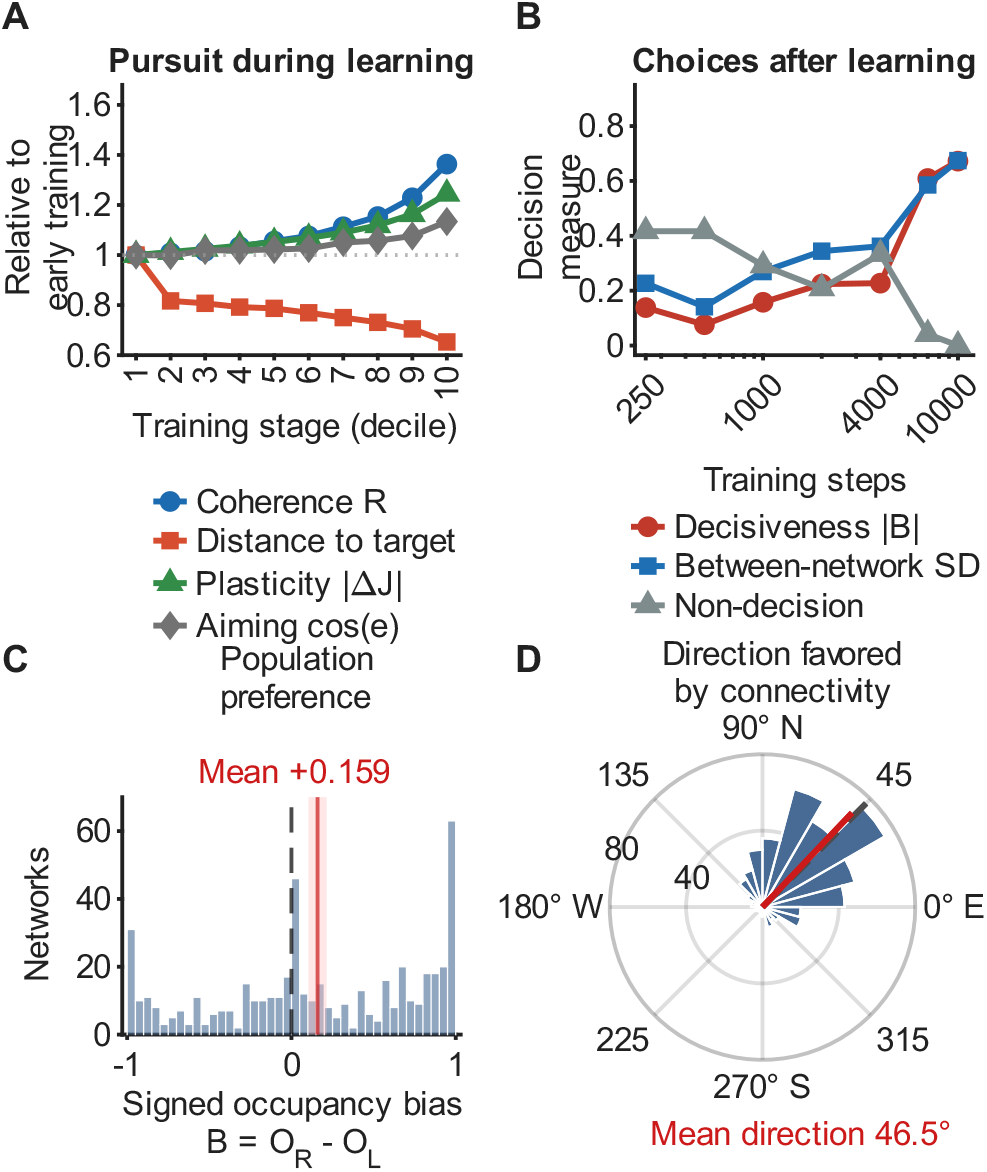
Learning experience shapes individual choices and population bias. (A) Skill acquisition over training. Directional coherence *R* rises and the agent’s mean distance to the target falls, while its instantaneous aiming error decreases modestly and perstep plasticity Δ*J* accelerates. Curves are population means over 500 networks across ten training stages. The dotted line marks unity, the early-training baseline. A training stage is one tenth of a single continuous training run; panel B uses snapshots of each of 24 independent training trajectories, so its axis shows steps rather than stages. (B) Decision-making develops with training experience. Connectivity is recorded at increasing training durations in each of 24 independent runs, then held fixed while the network is tested in the two-target choice; both decisiveness (the magnitude of the occupancy bias |*B* |) and between-network variation in signed bias rise with experience, while the fraction of runs that end without a decision falls to near zero. Committed, individually distinct decisions thus emerge over several thousand movement steps. (C) The population has a net rightward bias. The per-network signed-bias distribution is broad and two-sided, with mass near both *±*1 extremes, a third concentration of near-zero networks at the center, and a modest rightward excess of +0.159 (95% interval 0.104–0.214) rather than a uniform shift. Red line, the population mean; pink band, its 95% interval; black dashed line, zero bias. (D) The learned weights favor a direction in the world frame. Wedge length is the number of networks whose favored direction, read from the row sums of *J*, falls in each 15^*°*^ bin, with radial tick labels giving counts. The circular mean of that direction is 46.5^*°*^, in the northeast quadrant (bootstrap 95% interval 41.5–51.3^*°*^), aligned with the fixed 45^*°*^ bearing from which every network was trained, marked by the dashed line. Panels C and D use all 500 networks, with intervals from 100,000 networkbootstrap resamples. A mirror-start retraining control confirms that the offset follows the training frame. Networks trained from the reflected geometry develop the opposite, leftward offset, while the mirror-symmetric component is consistent with zero. Withintraining repeatability and the allocentric learning-rate sweep are shown in Supplementary Fig. S13.

#### 2. Training conditions shape individual preferences and population bias

Up to *l*_*H*_ = 10^*−*3^, higher learning rates increased the asymmetry and overall strength of the learned connections under the standard allocentric protocol. Tracking during training changed little across the lower rates, improved at *l*_*H*_ = 10^*−*3^, and failed at the highest rate as the weights saturated (Supplementary Fig. S13B). When tested with fixed weights, networks with stronger preferences could perform worse at both pursuit and choosing the stronger target, even before saturation (Supplementary Table S16).

Preference strength peaked near a training inverse noise of *β*_*H*_ = 14, exceeding that at the standard value of 17 by 0.154 across 100 matched seeds (95% interval 0.065–0.240), with no detected change in balanced accuracy (Supplementary Note S12). The standard training condition therefore produces individuality without maximizing its strength. Across 500 networks trained under the same protocol, greater summed connectivity change at the retained training steps was associated with weaker final bias (*r* = −0.338, *P* = 7.4 × 10^*−*15^). This correlation alone does not show that greater weight change weakens preference (Supplementary Note S7).

The initial training geometry also shaped population bias. Although individual networks preferred either side, their mean bias was rightward at +0.159 (Fig. 7C). The direction favored by the learned connections, estimated from their row sums, averaged 46.5^*°*^, close to the target’s initial bearing of 45^*°*^ (Fig. 7D). Mirroring the initial geometry reversed the population bias while preserving substantial differences among networks (Supplementary Note S3).

#### 3. Spatial reference frame shapes learned preferences and decision accuracy

To test whether learned individuality depends on how the network represents direction, we compared the allocentric model with an egocentric variant in which neurons’ preferred directions rotate with the agent’s heading. We trained 40 pairs of networks using matched random seeds, with each network retaining the same reference frame during training and testing (Supplementary Note S15; Supplementary Fig. S1). Across the five tested noise levels, *β* = 10, 16, 20, 40, and 60, allocentric networks showed substantial repeatability of signed bias, with ICCs ranging from 0.468 to 0.982. In egocentric networks, every ICC confidence interval included zero. This lack of detectable repeatability occurred despite considerable connectivity heterogeneity: the mean residual fraction was 0.117 in egocentric networks, compared with 0.087 in allocentric networks. The two models showed similar responses to learning rate during training, but their performance differed when the learned connections were frozen. Egocentric networks achieved higher orientation-balanced accuracy and lower mean pursuit error, although the confidence interval for the pursuit difference included zero at *β* = 10. At *β* = 40 and 60, their accuracy also exceeded that of their own gain-matched circulant controls. Thus, learned heterogeneity could improve choice accuracy without producing detectable repeatable target-side preferences. Its behavioral consequences depended on the spatial reference frame used during learning and testing.

### I. Continued plasticity limits the persistence of bias

The post-training behavioral tests above used fixed connection weights. To test whether individual preferences persist when experience continues to reshape the network (continued plasticity), we kept Hebbian learning active during the choice task at a fraction of the training rate.

At one percent of the training rate and without forgetting, preference strength remained high at 0.59 after 2,000 movement steps, but fell to 0.40 after 4,000 steps and 0.07 after 8,000. Faster learning changed the connectivity and weakened preferences more rapidly (Fig. 8A–E). Forgetting altered this outcome. When sufficiently strong relative to learning, it preserved preference strength over the tested interval and, at an intermediate learning rate, increased it (Fig. 8F).

**FIG. 8.**
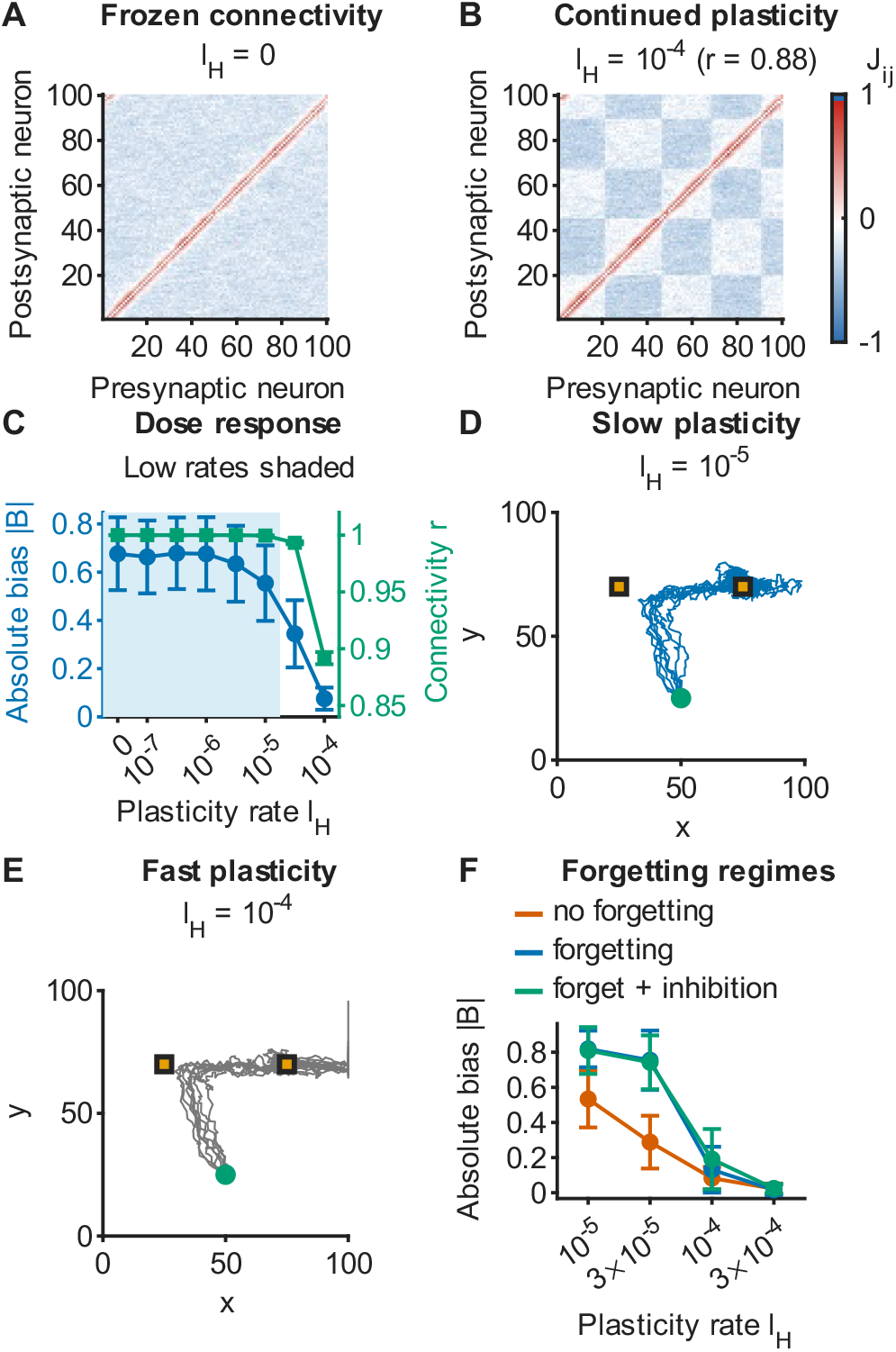
Over the tested interval, bias magnitude persists under continued plasticity that is slow relative to the original training. (A) Frozen learned connectivity of a representative network, a structured asymmetric ring. (B) The same connectivity after high-rate continued plasticity (*l*_*H*_ = 10^*−*4^); the ring is remodeled, and its correlation with the frozen ring is shown by the paired matrix comparison. Both are shown on the same diverging color scale. (C) Dose-response: the absolute equal-choice bias |*B*| (left) and the connectivity correlation *r* (right) against the continued rate *l*_*H*_ ; connectivity correlation remains high across the shaded low-rate window, while bias declines as *l*_*H*_ increases (20 networks). (D) Equal-choice arena trajectories of the representative network under slow continued plasticity (*l*_*H*_ = 10^*−*5^); the network commits to the side it prefers. In D and E the orange squares mark the two equal targets and the green circle the shared starting position. (E) Under high-rate continued plasticity (*l*_*H*_ = 10^*−*4^) the same network’s trajectories lose commitment and wander, ranging over the corridor between the targets and beyond both of them to the arena walls. (F) Continued plasticity with forgetting (*f*_*H*_ = 10^*−*3^) across four continued rates. While forgetting stays strong relative to the continued rate, up to *l*_*H*_ = 3 *×* 10^*−*5^, bias magnitude is preserved over the tested interval and at that rate rises to |*B* |*≈*0.74–0.75, whereas without forgetting it falls to 0.29. From *l*_*H*_ = −10^*−*4^ upward the preserved bias is smaller at the plotted *f*_*H*_ = 10^*−*3^ and bias magnitude declines in all three conditions, so the outcome depends on the ratio of forgetting to learning rather than on forgetting as such. Three conditions are shown: no forgetting, forgetting, and forgetting with the learning-time inhibition *I*_*H*_ = 10^*−*5^, the legend’s “forget + inhib”, which tracks the forgetting curve closely at every rate. Error bars in C and F show *±* 1.96 network standard errors. These population measurements establish persistence of bias magnitude, not retention of the same preferred side. See Supplementary Note S13.

Individual preferences can therefore persist during continued learning, but their strength depends on how quickly and for how long the weights change, and on the balance between learning and forgetting. Preserving preference strength does not establish that an agent retains its original preferred side, or that its preference remains stable indefinitely. Further analyses of connectivity, pursuit, choice accuracy, forgetting, and input offset are reported in Supplementary Note S13.

### J. Learning outcomes depend on initial connectivity

In the preceding experiments, networks began without recurrent connections, so learning built both the shared ring structure and the deviations associated with individual preferences. We next asked whether learning could generate similar preferences in a network that already had an organized, symmetric ring. This comparison tests whether individuality depends on how the connectivity develops.

Across 16 paired seeds, networks starting from zero developed strong preferences, with a mean absolute equal-choice bias of about 0.67, whereas the mean for those starting from the ring was 0.000 (Fig. 9). Without forgetting, the ring’s weights saturated. Forgetting prevented saturation but produced no consistent side preference.

**FIG. 9.**
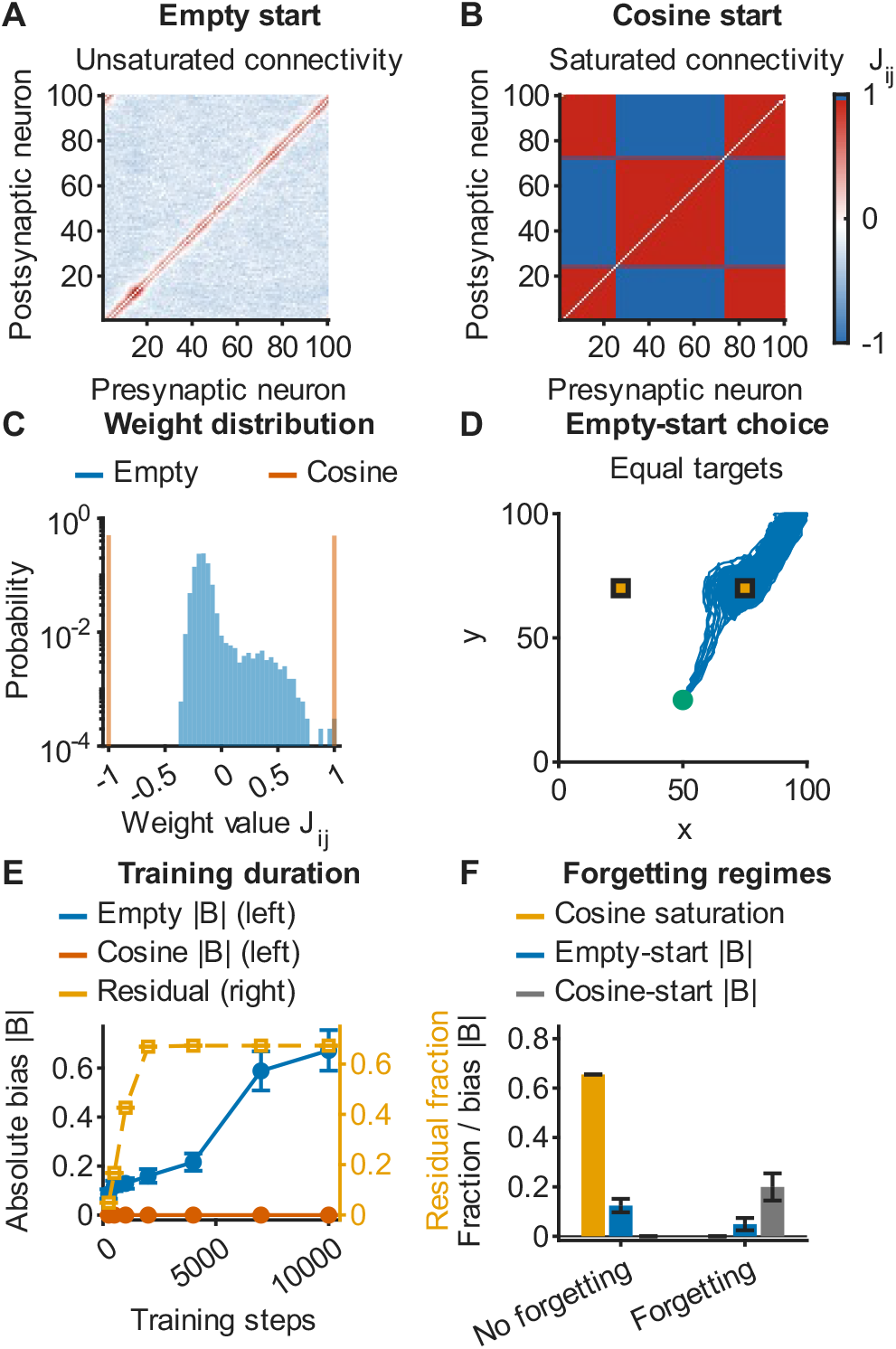
Individuality depends on the starting circuit; under the same learning a preformed ring saturates instead. (A)Connectivity trained from an empty start under standard Hebbian learning (*l*_*H*_ = 10^*−*3^, matched seed): a structured asymmetric ring with no saturated weights. (B) Connectivity trained from the symmetric cosine ring under the same learning, where plasticity without stabilizing decay drives the weights to their *±* 1 bounds, a saturated fraction of 1.0, giving saturated symmetric connectivity. Both are shown on the same diverging color scale. (C) Distribution of weight values. The empty-start weights spread around zero, while the cosine-start weights collapse onto*±* 1. (D) Equal-choice arena trajectories of the empty-start network, which commits to the target on its preferred side. (E) Absolute equal-choice bias against training length for the two starts, with the cosine start’s connectivity residual fraction on the right axis. The empty start’s bias rises overall to 0.67 while the cosine start’s stays at zero even as its residual fraction climbs to two-thirds, a large but symmetric residual (16 seeds). (F) Adding forgetting removes the cosine-start saturation but does not rescue the choice. This panel uses *l*_*H*_ = 10^*−*4^, a lower rate than the 10^*−*3^ of panels A–C, which is why its no-forgetting saturated fraction is 0.65 rather than the 1.00 of panel B. Three bar series share one axis: the cosine-start saturated fraction, which falls from 0.65 to zero once forgetting is added, and the absolute equalchoice bias of the empty start and of the cosine start in the same two conditions. With forgetting the connectivity no longer saturates, yet both starts remain weakly biased, with different mean magnitudes across the starts; these small cohorts do not establish equivalence. Panel F uses two independently trained seeds without forgetting and six with forgetting; error bars show*±* 1 network SEM. See Supplementary Note S12.

We then tested three other possible explanations. We rescaled the initial ring to match the learned networks’ overall coupling strength, varied training noise across the rescaled ring’s own transition, and added initial heterogeneity larger than the ring’s weight amplitude to break its symmetry. None of these changes produced preferences under the tested conditions, while the paired networks starting from zero developed preferences in every comparison. The outcome therefore depends on the initial connectivity, although these tests do not establish that an initially unstructured circuit is necessary for individuality. Further comparisons of learning rate, forgetting, connectivity changes, and choice accuracy are reported in Supplementary Note S12 and Supplementary Fig. S14.

## III. DISCUSSION

Hebbian learning can organize the circuit that controls movement and, in the same process, generate stable individual differences in spatial choice. Starting from neurons with fixed directional tuning and no recurrent connections, learning produced network-specific center–surround backbones together with smaller deviations from each network’s own average profile. Those deviations supported repeatable preferences when agents chose between equally attractive targets (Fig. 3). A shared learning rule and a shared training environment can therefore produce a common circuit architecture while allowing individuals to develop persistent behavioral differences.

Earlier work established that associative plasticity can organize recurrent circuitry for head-direction and spatial coding [34, 35]. Our results connect this process of circuit formation to individual preferences and their consequences across spatial tasks. They complement work on learning-dependent individuality [10] by identifying a specific recurrent-connectivity mechanism and testing it through matched interventions on the learned weights. This gives a mechanistic counterpart to evidence tying developmental variation in wiring to behavioral individuality [6, 40], and extends accounts of spatial choice built on geometry-dependent neural dynamics [55] by allowing the underlying connectivity to differ among individuals. Prescribed variation in connection weights and directional tuning also produced repeatable preferences. Learning is therefore one way to generate the heterogeneity underlying individuality, but repeatable preferences can also emerge from prescribed heterogeneity without learning.

The connectivity interventions show that commitment and flexible pursuit can coexist. Removing the residual and leaving each network’s mean ring profile strongly reduced side-preference strength and left little repeatability. Rearranging the residual while preserving its amount, the overall coupling gain and the mean and spread of the weights at every angular offset retained strong, repeatable preferences and markedly improved pursuit (Fig. 4; Supplementary Note S10). Within the tested networks, poor pursuit therefore depended on the particular arrangement acquired during learning and could be relieved without sacrificing commitment. These results provide no evidence for an obligatory trade-off between commitment and flexibility. Instead, the behavioral consequences of heterogeneity depend on its organization and the demands of the task. This is consistent with work on stochastic neural fields, where spatial heterogeneity pins a travelling bump and slows its diffusion by an amount set by its pattern as well as its magnitude [76].

Bias also has to be distinguished from decision quality. Its direction predicted whether a network did better with the stronger target on the left or on the right, whereas its magnitude showed no robust association with success averaged over both placements in the *β* = 60 cohort (Fig. 5). At some higher noise levels, stronger preferences were associated with lower accuracy, showing that the relationship depends on neural noise (Supplementary Table S41). Low accuracy repeatability and restricted variation limit interpretation of the high-*β* null. In the allocentric comparisons, heterogeneity narrowed the range of neural noise within which agents accurately chose the stronger target, without a consistent mean-performance advantage over the homogeneous reference (Fig. 6). Sensory input width also affected accuracy. At low neural noise, widening the standard input field eightfold raised orientation-balanced accuracy from 0.51 to 0.78, showing that poor performance partly reflected the narrow input rather than an unavoidable cost of learned individuality (Supplementary Table S26). Accuracy must nevertheless be interpreted alongside responsiveness. Because runs in which neither target is reached count as failures, the score combines reaching a target with selecting the stronger one. A value near 0.5 alone therefore does not establish insensitivity to target strength. Empirical work on lateralization and performance faces the same need to distinguish preference direction, preference strength, and failure to respond [14–17].

Examining both allocentric and egocentric representations reveals a further dependence on how the circuit controls behavior (Section II H 3). The egocentric networks remained heterogeneous, yet lacked detected repeatable target-side preference and achieved higher balanced accuracy, including an advantage over their own matched circulant controls at *β* = 40 and 60. This result shows that learned heterogeneity can improve accuracy even when repeatable target-side preferences are not detected. Because the reference frame changes in training and testing together, the comparison does not isolate effects on learning from effects on behavioral readout. The absence of detectable target-side repeatability under these conditions does not exclude other individual differences or repeatable preferences under other egocentric conditions.

The outcome depended on both the starting circuit and the experience that shaped it. Strong preferences developed from networks without recurrent connections, but not from a preformed cosine ring under the protocols we tested (Fig. 9; Supplementary Note S12). This demonstrates sensitivity to initial connectivity without establishing that individuality requires an initially unstructured circuit. Across networks trained under the same protocol, greater summed change at the retained training steps was associated with weaker final bias, *r* = −0.338, but this correlation does not show that greater weight change causes weaker preferences. Training geometry influenced the population distribution of preferences. The direction favored by the learned weights, estimated from their row sums, aligned with the initial target bearing. Mirroring that bearing reversed mean population bias from +0.153 to − 0.194 across 500 paired seeds trained at the main ensemble’s training inverse noise. Half the difference between the two population means was +0.174, with a 95% interval of 0.136–0.212 (Supplementary Note S3). Training orientation thus shifted the distribution of preferences without determining every individual’s preferred side.

Continued plasticity revealed limits to the persistence of preference strength. At one percent of the training rate, bias remained substantial through the standard test but declined with longer exposure, from 0.59 after 2,000 movement steps to 0.07 by 8,000. At the same learning rate, mean pursuit error increased from 16.3 to 24.8 over the standard test (Supplementary Note S13). Forgetting preserved preference strength, and sometimes increased it, when sufficiently strong relative to ongoing learning. Persistence therefore depended on rate, duration and forgetting. These measurements establish persistence of bias magnitude over finite intervals; they do not establish indefinite stability or continued preference for the same side.

Uniform global inhibition provided another way to alter behavior. Near the learned networks’ peak in spontaneous directional fluctuations, subtracting the same fixed offset from each neuron’s field strengthened preferences at moderate levels but suppressed responses to equal targets at larger levels (Supplementary Note S4). Together with the effects of neural noise and input width, this shows that the expression of individuality depends on the conditions under which the circuit operates.

The simplicity that makes the model tractable also limits its scope. The model represents a single angular variable through directional tuning. In its allocentric form, the resulting preference concerns a direction in the world rather than a side of the body, as in behavioral lateralization. These results concern the Hebbian update rule used here. Our interventions do not separate the effects of reciprocal asymmetry from other features of the learned heterogeneity. Behavior was also tested in a restricted set of arenas and target configurations. The standard input field has a standard deviation of one preferred-direction spacing, concentrating input on a few nearby neurons; we tested alternatives only by widening the field and changing target intensity. The structured angular variation in the learned weights should not be equated with twodimensional grid-cell firing. The shuffle comparison does not uniquely distinguish quasi-periodicity from smooth, nonperiodic structure. The learned networks are asymmetric and require a nonequilibrium description, so equilibrium arguments that apply to the symmetric reference do not transfer to them. Their dynamics also approximate an attractor rather than forming an exact continuous family of fixed points. Irregular experience produces quasi-attractor regimes, with limits on the variability those regimes can tolerate [77]. Extending the model to other sensory representations, plasticity rules and task geometries would test the generality of this mechanism. These results identify a possible circuit mechanism for individuality and suggest how to assess its behavioral consequences. Performance should be measured with the better option on either side of an individual’s preferred direction, and preference persistence should be assessed while allowing connectivity to continue changing. A shared learning rule can thus build a common circuit architecture while generating repeatable individual preferences; how those preferences affect behavior depends on the circuit’s organization, its spatial reference frame, and the task.

## IV. METHODS

### A. Neural and movement dynamics

An agent is controlled by a network of *N* binary neurons with states *s*_*i*_∈ {− 1, +1}. In the allocentric model, neuron *i* has a world-fixed preferred direction *α*_*i*_ = 2*π*(*i*− 1)*/N* and responds to stimuli according to the angular deviation between its preferred direction and the stimulus bearing. Circular angular distance is *δ*(*a, b*) = min{| *a*− *b* |, 2*π*−| *a*− *b*|. A stimulus *t* at bearing *θ*_*t*_ contributes a circular Gaussian input to each neuron,

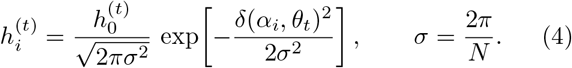

Bearings are computed with the two-argument arctangent, *θ*_*t*_ = atan2(*y*_*t*_− *y, x*_*t*_ −*x*), wrapped to [0, 2*π*). A target coincident with the agent supplies no directional stimulus. A zero population vector produces no displacement and retains the previous orientation. A global inhibition *h*_*b*_ is subtracted once from the field of each neuron, independently of the number of stimuli. Raising it lowers the drive reaching every neuron, so it sets how strongly the network responds during testing. It represents a constant inhibitory background of the kind that balances excitation in real circuits [19, 20]. The incoming local field of neuron *i* is

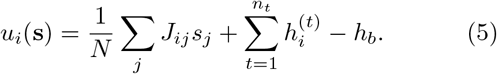

Neural states evolve by a single-spin Metropolis-like local flip rule. At each neural update one neuron is chosen uniformly at random and its flip *s*_*i*_→ − *s*_*i*_ is accepted with probability

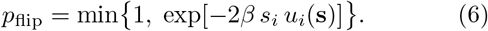

When the connectivity is symmetric, Eq. 6 is the Metropolis rule for the equilibrium Hamiltonian

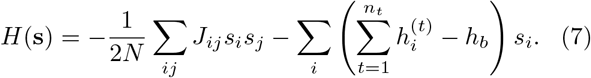

Learned networks are in general asymmetric and have a zero diagonal, so no scalar energy governs their dynamics; they are treated as nonequilibrium stochastic networks defined directly by their local fields, Eq. 5. For those networks *β* is an inverse noise, or equivalently a multiplicative factor on the local field, and not an inverse temperature, since there is no Hamiltonian for the dynamics to equilibrate to. We use the thermodynamic interpretation only for the symmetric cosine reference, where Eq. 7 applies.

Unless stated otherwise *J*_*ii*_ = 0 and *h*_*b*_ = 0. The coupling sum has the conventional 1*/N* normalization, so the energy is extensive and *β, h*_0_, and *h*_*b*_ are the physical inverse noise, stimulus amplitude and input offset reported on all axes; the equivalent parameters for an unnormalized recurrent sum are related by *β* = *Nβ*_sim_, *h*_0_ = *h*_0,sim_*/N* and *h*_*b*_ = *h*_*b*,sim_*/N*. Equation 7 is the total energy of a network in these units, so an energy per neuron is *H/N* and a heat capacity per neuron is *β*^2^ Var(*H*)*/N*. The stimulus-free transition diagnostics use this convention throughout.

The agent moves along the vector sum of the preferred directions of its active neurons. After *T*_0_ = 50*N* neural updates its position advances by

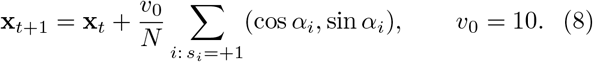

In the egocentric variant, the world direction of neuron *i* is instead *α*_*i*_(*t*) = 2*π*(*i*− 1)*/N* + *ϕ*(*t*), modulo 2*π*, where *ϕ*(*t*) is the agent’s heading at the start of the movement step. These current directions are used in both the sensory input of Eq. 4 and the movement readout of Eq. 8. After decoding movement, *ϕ* is set to the population-vector heading for the next step. Connectivity remains attached to neuron identities, and each condition uses the same reference frame during training and testing (Supplementary Note S15).

In the behavioral probes a moving target updates each coordinate independently by *v*_*t*_ sign(*z*) with *z*∼ *N* (0, 1), followed by the arena boundary operation, so *v*_*t*_ is a per-coordinate random-walk step and the Euclidean displacement away from boundaries is 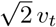; the training task uses a Gaussian step of the same scale, given below. Training and two-target choice clamp each coordinate to [0, *L*]. Standard pursuit and spontaneous-motion probes instead use periodic boundaries, wrapping coordinates modulo *L* and using minimum-image target bearings and distances. The continued-plasticity pursuit comparison uses fixed boundaries at all learning rates, including its frozen reference (Supplementary Note S13). Neural states are initialized independently and uniformly at random on{−1, +1} at the start of every run, so no run starts with an activity bump left over from the previous run.

The neural-update count *T*_0_ = 50*N* was fixed by comparison with a 100*N* reference. Across mean absolute bias, target-zone occupancy, and orientation-balanced accuracy, the largest condition-mean deviation from the reference was 0.073 at 20*N* and 0.050 at 50*N*, both within the tolerance of 0.08; we use 50*N* throughout the behavioral probes (Supplementary Fig. S15). A moving target uses one bearing channel and a two-target choice uses two. Further numerical detail and the local-field validation are given in Supplementary Note S1.

### B. Connectivity

The homogeneous reference network uses the cosine profile widely adopted for continuous attractor models,

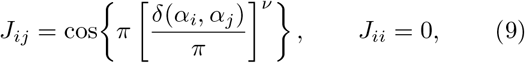

with tuning exponent *ν*; the reference used in the individuality comparisons has *ν* = 0.5. For *ν* = 1 this is a cosine of the circular distance. The coupling changes sign at an angular separation of *π*2^*−*1*/ν*^, which is 90^*°*^ at *ν* = 1 and 45^*°*^ at the *ν* = 0.5 used in the main comparisons, so the profile used here is still center–surround but has a narrower excitatory core and a broader inhibitory surround than a plain cosine. We continue to call it the cosine ring for brevity.

We call *J* the connectivity and its entries weights. A weight is not a model of one synapse. A single *J*_*ij*_ may take either sign and may change sign during learning, so it stands for the net influence one neuron has on another rather than for a contact with a fixed excitatory or inhibitory identity. By heterogeneity in the weights we mean differences between these net influences.

As a second, independent source of connectivity heterogeneity we also use prescribed networks. Random noise is added in two constructions that serve different purposes. The *clipped-uniform* random-noise networks add to each weight of the *ν* = 0.5 ring a value drawn uniformly on [−1, 1], subject to *J*_*ij*_ remaining in [−1, 1]; their overall coupling scale is left free, and they are the secondary comparison in Fig. 5. The *matched-gain* randomnoise networks instead add independent Gaussian perturbations of standard deviation 0.3 to a ring already scaled to the reference gain of 19.6, then rescale the perturbed matrix to that same gain; these are used in comparisons with normalized reference networks, including Fig. 6. The matched-gain tuning networks assign neuron-specific tuning exponents *ν*_*i*_ = *ν*_0_ + *σ*_*ν*_ *η* with *ν*_0_ = 0.5, *σ*_*ν*_ = 0.3 and *η* standard normal, clipped to *ν*_*i*_ ∈ [0.1, 2], and are rescaled to the same Frobenius norm. The reference gain is fixed at 19.6; learned networks remain unscaled, whereas paired intervention controls match each learned network’s actual gain. The prescribed randomnoise networks provide a generality check on the learned results and are secondary to them, so the central connectivity interventions act within each learned connectivity rather than between one network type and another.

For any connectivity *J* the directed lag is *d* = (*j*− *i*)mod *N*, which counts how far apart two neurons lie around the ring of preferred directions. The network-specific circulant profile *C* assigns to every entry at lag *d* the mean across rows of that same network at that lag. Thus *C* is shared across neurons within one network, but it is not a population-average profile; the residual *R* = *J* − *C* contains the neuron-specific departures. Normalized asymmetry is ∥*J* − *J*^T^ ∥_*F*_ */*∥ *J* ∥_*F*_ and the antisymmetric part of a matrix is (*J*− *J*^T^)*/*2. Averaging over rows at a fixed lag commutes with transposition, so *C* and *R* each have their own antisymmetric part, and the two sum to the antisymmetric part of *J*. We therefore measure the two separately instead of inferring either from the value for the whole matrix (Supplementary Note S7). Residual spectral concentration is computed from the nonzero-frequency power of the lag-ordered residual after excluding the protected local region. An exact longrange shuffle independently permutes the long-range entries within each row without replacement, preserving both the protected local entries and the row-wise weight multiset. The matched replacement *Q* used in the dose 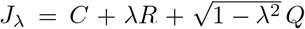is built one lag at a time. Within each directed lag *d* the entries of *Q* are drawn independently from a normal distribution and centered to zero mean, which is also the mean of *R* at every lag, since *R* is a residual. They are then made orthogonal to *R* at that lag by removing their component along it, and rescaled to the norm of *R* at that lag. The diagonal is lag *d* = 0, where *R* vanishes and so does *Q*. Lag means, lag variances, and residual norms ∥*Q*∥ _*F*_ = ∥*R* ∥_*F*_ are matched exactly, with *Q* orthogonal to *R* within every lag. Together with lag centering, these conditions preserve the full-network norm, ∥*J*_*λ*_ ∥_*F*_ = ∥*J*∥ _*F*_, in the dose of Eq. 12; realized errors are below 10^*−*12^ (Supplementary Note S10). The dose therefore keeps the average weight and the spread of weights at every separation, and hence the overall coupling strength, while replacing the arrangement of those weights.

### C. Hebbian learning

During training, plasticity is switched on. Learning uses no externally supplied error signal or reward. Every training run starts from an empty recurrent connectivity, *J*_*ij*_ = 0, except in the preformed-ring study of Section II J, which starts each run from the *ν* = 0.5 cosine ring of Eq. 9. The prescribed networks above are never trained. Their heterogeneity is set by hand and then held fixed, which makes them a control for the learned networks.

#### a. The training task

Pursuit of a single moving target is the only task on which any network is trained. The agent starts at the corner (0, 0) of a fixed-boundary arena of side *L* = 30 and the target starts at the arena center (15, 15), so on the first movement step the target lies at a 45^*°*^ bearing and a separation of 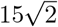. Thereafter the target random-walks freely, each coordinate stepping independently by a Gaussian draw of standard deviation *v*_*t*_ = 1 per movement step, followed by the same boundary clamp as the agent, and the agent pursues it for 10,000 movement steps while its weights change under Eq. 10. The drive of Eq. 4 is a function of the target’s bearing alone and its amplitude *h*_0_ is fixed, so approaching the target neither recruits additional neurons nor drives them harder; proximity instead changes how fast the bearing turns. A stimulus that decays with distance is used only in the homogeneous-agent check of Supplementary Note S5.

#### b. The plasticity rule

The correlations the rule acts on are generated by the task and by the topographic layout of the input, not imposed. A target at a given bearing drives the contiguous group of neurons tuned near it into the active state together while neurons tuned farther away receive little direct drive, so over a pursuit run pairs with similar preferred directions are repeatedly in the same state and pairs with distant preferred directions are not. Once the flip of Eq. 6 has been accepted or rejected, the row of the selected neuron *i* is updated from the resulting post-flip network state,

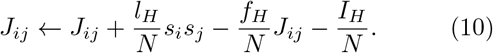

Here *l*_*H*_ is the learning rate, *f*_*H*_ the forgetting rate, and *I*_*H*_ the learning-time inhibition, also called the weight drift, a constant negative term acting on the weight itself rather than on any neuron; all three have the same 1*/N* normalization as the recurrent sum of Eq. 5. The first term is the product of two ± 1 neural states, so neurons in the same state, whether both active or both inactive, increase their coupling weight, and neurons in opposite states decrease it [47, 48]. Its mean contribution is proportional to the mean spin product at the times the row is updated. We therefore call it a spin-correlation rather than a coactivity rule. The second term is the forgetting, which decays each weight in proportion to its own magnitude. Immediately after the update the diagonal entry *J*_*ii*_ is reset to zero and the whole row is clipped to [1, − 1]. The update is applied at each of the *T*_0_ neural updates within a movement step, so plasticity and the fast neural dynamics interleave rather than run in separate phases. The target bearing, and with it the drive of Eq. 4, is refreshed once per movement step and held fixed across those *T*_0_ updates. Pure spin-correlation learning is recovered by setting *f*_*H*_ = *I*_*H*_ = 0.

#### c. The standard training task

The *standard training task* is the protocol just described, run with *N* = 100, *β*_*H*_ = 17, *l*_*H*_ = 0.001, *f*_*H*_ = 0, *I*_*H*_ = 10^*−*5^, *T*_0_ = 100*N* neural updates per movement step, target coordinate-step standard deviation *v*_*t*_ = 1, receptive-field strength *h*_0_ = 0.02, arena side *L* = 30, 10,000 movement steps, and zero input offset. The learned networks analyzed throughout are 500 independent realizations of it that share all of these parameters and differ only in their random seed. Other training protocols vary named parameters of this one and are stated with the figure that uses them.

#### d. The starting geometry and the shared directional bias

The starting geometry is deliberately the same for every network. Holding it fixed means the 500 networks differ only in the stochastic walk of their target and in their own neural noise, so the differences that accumulate in their connectivities arise under a shared protocol rather than from different initial viewing angles. This design does not separate the contributions of target-path variation, neural noise, and their feedback through movement. The drawback is that the shared first step leaves a common direction in the trained population, which is therefore not perfectly symmetric. The population has a modest rightward occupancy offset, and the direction its learned weights favor, taken as the angle of the first circular Fourier mode of the row sums of *J*, averages 46.45^*°*^ in the world frame, close to the fixed 45^*°*^ starting orientation. This common direction, and the side preference that goes with it, make up the shared directional bias described in Section II A. The original population distribution and connectivity direction are shown in Fig. 7C,D; the paired mirror-start retraining test that attributes the offset to the training frame is reported in Supplementary Note S3.

#### e. Testing with frozen connectivity

No network encounters a choice between targets during initial training. Post-training behavioral tests use frozen connectivity, except in the continued-plasticity study of Section II I. In that study Eq. 10 continues to act during behavior, at a learning rate well below the training value.

### D. Behavioral tasks and outcomes

Choice tasks use *N* = 100 neurons in an arena of side *L* = 100 with fixed boundaries, two targets at (25, 70) and (75, 70), the agent starting at (50, 25), a target-zone radius of 10, and 2,000 movement steps. Equal-choice targets have intensity *h*_0_ = 0.01. Unequal choices use intensities (0.01, 0.015) and (0.015, 0.01) in matched orientations so that a network is tested with the stronger target on each side. Standard pursuit uses a single moving target over 1,500 steps with *v*_*t*_ = 1 and periodic boundaries. The first 200 movement steps are excluded from occupancy, distance, speed, and tracking summaries but retained for first-entry and first-choice outcomes.

Signed occupancy bias is *B* = *O*_*R*_ − *O*_*L*_, where *O*_*R*_ and *O*_*L*_ are the post-transient fractions of observations inside the right and left target zones; it is zero when neither zone is occupied. A network’s signed bias is the mean of *B* across its nested repeats, and its primary bias magnitude is the absolute value of that mean, 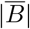, rather than the mean of trialwise| *B*|.

The primary decision measure is orientation-balanced first-entry accuracy. Within each network, first-entry accuracy is averaged across the matched right-stronger and left-stronger tasks, and a trajectory that reaches neither target is scored as an incorrect non-decision. Balancing orientation in this way prevents a stable side preference from being read as a general decision advantage whenever it happens to align with the stronger option. In the primary repeated-trial cohort, Hebbian networks are evaluated at *β* = 40 and 60 and prescribed random-noise networks at *β* = 16, and every accuracy statement is reported at the *β* at which it was measured.

Two occupancy-conditional ratios measure how exclusively a network commits to one target:

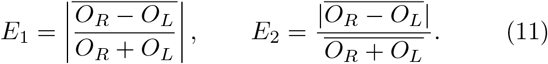

The overbar denotes the average across a network’s nested repeats. For *E*_1_, the per-trial ratio is set to zero when *O*_*R*_ + *O*_*L*_ = 0; *E*_2_ is set to zero when the mean occupancy 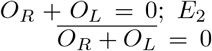. Both ratios can saturate at one even when occupancy is sparse, so their associations are reported with rank correlations, explicit occupancy adjustment, saturation diagnostics, and held-out prediction.

Choice-visit consistency is the fraction of target-zone observations, from first entry onward, that fall at the first-selected target, with observations outside both zones excluded from the denominator. First passage is the first complete-trajectory entry into either zone, capped at the trajectory length for latency comparisons. Tracking error is the post-transient mean distance from the agent before its movement to the target after its movement, and tracking performance is its negative. Distances measured before target movement and after agent movement, and paired probes with Gaussian instead of diagonal target increments, are reported as sensitivity controls. These conventions are collected in Supplementary Table S19, and extended motivation is given in Supplementary Note S9.

### E. Connectivity interventions

To ask whether the particular learned arrangement, rather than the presence of heterogeneity, drives behavior, each learned connectivity is compared with controls that hold its lower-order connectivity statistics fixed. Writing *J* = *C* + *R*, a moment-matched dose series is

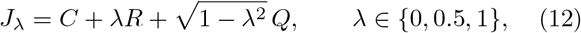

where *Q* is zero-mean, orthogonal to *R*, and matched in norm within every directed lag, so that lag means, lag variances, and whole-connectivity Frobenius gain are preserved while the amplitude of the learned residual varies.

The *reshuffled* control keeps, at each directed lag, the same list of departures from *C* and assigns them to different pairs of neurons, so only their positions change. The *wave-shifted* control keeps the size of every Fourier component of the residual at each lag and moves its peaks elsewhere around the ring. Because removing the residual entirely also lowers total coupling gain, a gain-matched circulant control scales each circulant profile to the exact Frobenius norm of its learned connectivity while retaining translational symmetry. Structural matching errors were below 10^*−*12^. The intervention analysis uses 100 learned networks at *β* = 60 with three nested repeats. Its outcomes are absolute equal-choice bias, choice-visit consistency, capped first-passage performance, orientation-balanced accuracy, and pursuit performance; learnedversus-control contrasts are Holm-adjusted across the set of tests they belong to. Network-mean neural flip rate is the fraction of neuron-time pairs that change state between consecutive retained samples. Full construction details are given in Supplementary Note S10.

### FSpatial concentration, entropy, and transition controls

Learned networks show multiple peaks in neural activity across movement directions, and a minority of them return repeatably to the same off-target site. Both are reported in full in Supplementary Note S8 with Supplementary Fig. S4. Off-target spatial concentration is scored with held-out repeats. Candidate off-target centers are selected from one set of repeats after excluding target neighborhoods, and their concentration is evaluated on a separate repeat, so that the same trajectory cannot both select and score a hotspot. A network is counted as showing a repeatable trap only when held-out off-target occupancy and repeat-to-repeat center location both meet fixed thresholds. Controls are the exact row-wise longrange permutation, a long-range-smoothed connectivity, and a gain-matched version of the smoothed connectivity; thresholds and the robustness grid are reported in Supplementary Note S8.

Neural entropy is computed from five independent neural-state repeats for each of 100 learned networks in equal choice, pursuit, and stimulus-free motion. For pursuit, the principal measure is the mean across neurons of binary activation entropy conditional on the presented target direction, *N*^*−*1^∑_*i*_ *H*(*A*_*i*_| *θ*), with a firstorder finite-sample correction; choice and stimulus-free entropy are marginal over direction. These quantities describe sampled single-neuron state variability, rather than a temporal entropy rate or the joint entropy of the population state. It is treated as a repeatable descriptor of neural variability whose behavioral associations are examined after adjustment and by held-out prediction (Supplementary Note S14).

Order–disorder transitions (Supplementary Fig. S12) are located in stimulus-free symmetric cosine networks at *N* ∈ {50, 100, 200} over a nonuniform *β* grid, using directional susceptibility, the Binder cumulant, and the integrated relaxation time of the magnitude of the first circular Fourier mode, cross-checked against a spectral linear-instability estimate. Asymmetric learned and random-noise networks are summarized with the same dynamical diagnostics but are not assigned equilibrium critical points. Input-offset modulation changes directional bias and target occupancy without changing the connectivity norm. The offset *h*_*b*_ is implemented once per neuron and varied separately for the learned and the random-noise networks. A sensitivity analysis compares this global offset with an offset applied per target channel: the total inhibition is doubled for two-target trials and unchanged for single-target pursuit. The sweeps in Supplementary Fig. S12E,F use *β* = 80 for the learned networks and *β* = 10 for the clipped-uniform randomnoise networks; corresponding quantitative comparisons at *β* = 60 and *β* = 16 are reported in Supplementary Note S4.

### G. Network classes and coupling measures

Four network types are compared in this paper. They are the symmetric cosine ring, the learned Hebbian networks, the prescribed random-noise networks, and networks given heterogeneous tuning. Selected fixed-noise comparisons are summarized in Table I; additional noise sweeps and the separate gain-matched Gaussian-noise controls are specified in Supplementary Note S4. We use *near-critical* only for the symmetric cosine ring near its order–disorder transition at *β*≈ 2.7. That transition is an equilibrium one, located by four independent stimulus-free diagnostics, three of which are plotted in Supplementary Fig. S12A–C. No single transition value is assigned to the learned networks. Their susceptibility peaks at a median *β* = 58 while the relaxation time of the directional-order magnitude peaks at a median *β* = 120 on the extended grid, so the two diagnostics do not agree on one value. Both are finite-duration measurements on networks that are asymmetric and out of equilibrium, so the two peaks describe what was measured and do not locate a critical point. Training uses the same inverse-noise parameter, at *β*_*H*_ = 17. Because connectivity changes during training, this value does not identify a corresponding point on the frozen-network response curves. Neither selected learned-network value is where accuracy is highest, which is near *β* = 16 (Section II G).

Three different quantities measure coupling strength in this paper, and they are not interchangeable. *Gain* always means the Frobenius norm ∥*J*∥ _*F*_, the quantity held at 19.6 in the prescribed reference classes and at each learned network’s own value in its matched intervention controls. *Scale factor* means a multiplicative constant applied to a whole connectivity matrix, as in the interpolations between learned and prescribed structure. *Spectral radius* means the largest eigenvalue modulus, and is used only as a saturation diagnostic, where a value approaching *N* signals collapse onto a single stored pattern. The number of neural updates per movement step, *T*_0_, differs by context and is collected in Table II. The behavioral value was fixed by convergence against a 100*N* reference (Supplementary Note S1).

**TABLE II.** Neural updates per movement step, *T*_0_, by context.

| Context | $T_0$ | Note |
| --- | --- | --- |
| Behavioral probes, all choice and tracking tasks | 50$N$ | converged against a 100$N$ reference |
| Training of the analyzed 500-network ensemble | 100$N$ | |
| Connectivity-emergence sweeps of Fig. 2A–C | 200$N$ | longer integration for the structural sweeps; the $N = 400$ sweeps of panels D and E use 100$N$ |

### H. Statistical analysis

Tracking error is reported for several subsets and protocols and its absolute value differs accordingly. The intervention set of 100 networks at *β* = 60 gives 13.0, the 40-network continued-plasticity set at the same *β*, with fixed boundaries, gives 16.3, and the training-time measurements in the small training arena are on a different scale again. Only contrasts computed within one protocol are compared. The absolute equal-choice bias 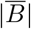 of the frozen learned networks likewise differs across subsets: it is 0.537 in the intervention cohort and 0.676 in the continued-plasticity dose cohort. These baselines should not be treated as paired comparisons.

The independently generated network is the unit of analysis. Stochastic trajectories are nested repeats and are averaged within network before population tests, except where the repeatability model retains them. Means, paired differences, and correlations are accompanied by 95% intervals from network-level bootstrap resampling, with 10,000 draws for behavior and interventions, 100,000 for the analysis of the shared directional bias in Fig. 7C,D, and 5,000 for entropy analyses. Repeatability is reported as the intraclass correlation, ICC, with networks as groups; its one-way analysis-of-variance estimator, which can take small negative values, is defined in Supplementary Note S9. Pearson correlations are accompanied by Spearman and leave-one-network-out sensitivities, and predictive claims use repeated ten-fold crossvalidation in which every scaling operation is fit on the training fold; the reported cross-validation range reflects variability across fold assignments rather than a population confidence interval.

Intervals are percentile intervals of the network-level bootstrap. When all observed networks have the same outcome, these empirical intervals have zero width; this does not imply certainty about the population probability. Unless another test is stated, paired contrasts use two-sided *t* tests on network-level differences. Pearson correlations use the statistic 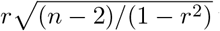 with *n* − 2 degrees of freedom; Spearman correlations use the corresponding rank test over the *n* networks. Partial correlations are tested on residualized variables, subtracting one further degree of freedom for each covariate. Two-sided *P* values are Holm-adjusted within each set of outcomes tested together, with correlations forming their own set. For common-random-number comparisons, matched conditions draw the same number of variates regardless of the active set. This preserves the intended coupling after their neural states diverge. Numerical settings are described in Supplementary Note S1.

#### a. Evolutionary dynamics

A population of *M* agents with heritable, variable *β* competes to find and follow a target, and more successful individuals replace others. Selection follows a pairwise comparison. At each step two agents, *i* and *j*, are drawn at random, and *j* replaces *i* with the Fermi probability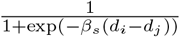,where *d*_*i*_ is the average distance of agent *i* from the target, the payoff is the negative of that distance, and *β*_*s*_ is the selection strength. Replacement is subject to mutation, which occurs with probability *µ*_1_ = 0.01. When a mutation occurs, the offspring value is *β*_*o*_ = *β*_*p*_ + *µ*_2_*η*, where *β*_*p*_ is the parent’s value and *η* is a standard normal draw. The mutation amplitude *µ*_2_ is set to *N/*10, where *N* is the number of neurons in the agent, so it differs between the two network sizes compared. Each population contains *M* = 100 agents; selection strength is *β*_*s*_ = 1. Initial physical *β* values are uniform on [0, *N*], and mutated values are clamped to this interval. A clone inherits its parent’s sampled fitness; every mutation triggers a new pursuit evaluation, including a mutation clamped at a boundary. We run ten independent populations for each network size and target speed for 10,000 pairwise updates, equivalent to 100 generations. This experiment uses the *ν* = 1 cosine ring, rather than the *ν* = 0.5 ring of the main comparisons. The evolutionary results are reported in Supplementary Note S6

### I. Simulation parameters by figure

All simulations use *N* = 100 binary neurons unless stated. The *standard choice arena* is a square of side *L* = 100 with fixed boundaries, equal targets at (25, 70) and (75, 70), agent start (50, 25), target-zone radius 10, *v*_0_ = 10, *T*_0_ = 50*N* neural updates per movement step, 2,000 movement steps with a 200-step transient, and equal-choice intensity *h*_0_ = 0.01; unequal choice uses matched intensities (0.01, 0.015) and (0.015, 0.01); standard single-target pursuit is one moving target (*v*_*t*_ = 1, *h*_0_ = 0.01) over 1,500 steps with periodic boundaries. The connectivity-intervention choice trials instead last 1,800 steps; this full horizon is both the first-entry censoring cap and the latency-performance denominator, while occupancy still excludes the first 200 steps. The *standard training task* is the protocol and the parameters set out above under *Hebbian learning*. Intervals are 95% network bootstraps with 10,000 draws unless stated.

#### a. Figure 2

Panels A–C use the connectivity-emergence training sweeps: standard training task geometry with *T*_0_ = 200*N*, learning rates *l*_*H*_ ∈ {0.001, 0.01, 0.1}, forgetting *f*_*H*_ = 0.001, weight drift *I*_*H*_ = 0.0001, over a 10,000 movement-step horizon; the training inverse noise *β*_*H*_ varies along the *x*-axis of A. Panels D and E show one *N* = 400 realization per plotted condition, at *l*_*H*_ = 0.01 with *f*_*H*_ = 0, *I*_*H*_ = 10^*−*4^ and *T*_0_ = 100*N*. The *β*_*H*_ = 12 condition uses *σ* = 8*π/N*, and the *β*_*H*_ = 39 condition uses *σ* = 2*π/N*. Panel F uses the 500 networks of the standard training task; residual spectral concentration excludes the protected local region (circular lag ≤ 7) and is compared to the exact long-range shuffle, with network-bootstrap intervals over the 500 networks.

#### b. Figure 3

Learned-network panels (A, C, D) are the equal-choice task at *β* = 40 in the standard choice arena, 100 networks with five repeats each. The homogeneous reference (B, D) is the cosine ring of Eq. 1 with *ν* = 0.5 at *β* = 4, 500 independently seeded trajectories. The same settings generate the trajectory examples of Supplementary Fig. S5.

#### c. Figure 4

All panels use *β* = 60 in the standard choice geometry with 1,800-step choice trials, 100 learned networks, and three nested repeats. The moment-matched dose is taken at *λ* ∈ {0, 0.5, 1} : connectivity profiles are shown in A and B, equal-choice outcomes in C and F, unequal-choice accuracy in D, and pursuit in E.

#### d. Figure 5

Standard choice arena; Hebbian networks at *β* = 60 (and *β* = 40 in D), random-noise networks at *β* = 16; 100 independently generated networks of each type with five nested repeats; right-/left-stronger accuracy scored on the complete trajectory. Randomnoise networks add to each weight a value drawn uniformly on [−1, 1] subject to *J*_*ij*_ ∈ [−1, 1].

#### e. Figure 6

Standard choice arena, with prescribed networks held at the reference Frobenius gain of 19.6 and learned networks left at their own gain and taken over *β*∈{4, 16, 20, 40, 60, 80, 30} networks of each type at each *β*. Balanced accuracy is the orientation-balanced first-entry accuracy on the 1.5:1 unequal choice, averaged over the matched right-stronger and left-stronger tasks; repeatability is the ICC of the signed equal-choice occupancy bias across five independent runs of each network. Panel C plots the two against each other for the learned networks alone. Error bars in B are ± 1.96 standard errors of the mean across the 30 networks of each type; the shaded band in A and B marks *β* = 16–20.

#### F. Figure 7

Training as in the standard training task. Panel A uses the 500-network ensemble, differing only by training seed, and splits each run into deciles (100 frames retained at ten-step intervals each). B follows 24 independent training realizations and freezes snapshots of each at seven training lengths from 250 to 10,000 movement steps, scoring the two-target choice at *β* = 60. Panels C and D use all 500 networks with 95% intervals from 100,000 network-bootstrap resamples; the signed bias is *B* = *O*_*R*_ − *O*_*L*_ (positive = rightward) and the direction of the shared bias is the angle of the first circular Fourier mode of each connectivity’s row-sum profile (0^*°*^ = east). Within-training repeatability and the allocentric learning-rate sweep are described in Supplementary Note S7 and Supplementary Fig. S13.

#### g. Figure 8

Continued plasticity applies the Hebbian correlation term after every neural update at a reduced rate *l*_*H*_ during behavior at *β* = 60 in the standard choice arena, with no forgetting or learning-time inhibition; the dose ladder spans *l*_*H*_ ∈ {0, 10^*−*7^, 3 × 10^*−*7^, 10^*−*6^, 3×10^*−*6^, 10^*−*5^, 3×10^*−*5^, 10^*−*4^} on 20 networks. Panel F compares runs without forgetting, with forgetting (*f*_*H*_ = 10^*−*3^), and with forgetting and inhibition, across *l*_*H*_ ∈ {10^*−*5^, 3× 10^*−*5^, 10^*−*4^, 3 ×10^*−*4^.

#### h. Figure 9

Sixteen seed-paired training runs per condition: empty start (*J* = 0) versus cosine start (*ν* = 0.5 ring) under the standard training task, with connectivity recorded at training lengths from 250 to 10,000 movement steps and probed at *β* = 60 in the standard choice arena. Panel F uses *l*_*H*_ = 10^*−*4^ both without forgetting (saturated fraction and biases from two seeds) and with forgetting *f*_*H*_ = 10^*−*3^ (six seeds).

## AUTHOR CONTRIBUTIONS

M.S.: conceptualization, methodology, software, validation, formal analysis, investigation, data curation, visualization, writing—original draft. I.D.C.: methodology, validation, funding acquisition, supervision, writing—review and editing. Both authors reviewed and approved the final text.

## CODE AND DATA AVAILABILITY

Simulation and analysis code, figure-generation inputs, and derived data are provided in Figshare [78] (DOI: 10.6084/m9.figshare.34307781; private review link).

## ACKNOWLEDGEMENTS

The authors acknowledge funding from the Deutsche Forschungsgemeinschaft (DFG, German Research Foundation) under Germany’s Excellence Strategy—EXC 2117–422037984; the DFG Gottfried Wilhelm Leibniz Prize 2022 584/22 (I.D.C.); the Max Planck Society; DFG project number 462886202; the European Union’s Horizon 2020 Research and Innovation Programme under the Marie Sk∤odowska-Curie grant agreement no. 860949; the Struktur-und Innovationsfonds für die Forschung of the State of Baden-Württemberg; the Pathfinder European Innovation Council Work Programme no. 101098722; and Office of Naval Research grant N0001419-1-2556.

We acknowledge the use of large language model assistants, OpenAI’s ChatGPT Codex (5.5, 5.6, 6) and Anthropic’s Claude Code (Opus 4.8 and 5), for assistance with plotting and analysis codes, readability suggestions and language edits, and critical examination of the scope of the claims. The authors reviewed and verified all analyses, calculations, results, and text and take full responsibility for the content of the paper.

